# Ribosome profiling at isoform level reveals an evolutionary conserved impact of differential splicing on the proteome

**DOI:** 10.1101/582031

**Authors:** Marina Reixachs-Solé, Jorge Ruiz-Orera, M Mar Albà, Eduardo Eyras

## Abstract

The differential production of transcript isoforms from gene loci is a key cellular mechanism. Yet, its impact in protein production remains an open question. Here, we describe ORQAS (ORF quantification pipeline for alternative splicing), a new pipeline for the translation quantification of individual transcript isoforms using ribosome-protected mRNA fragments (Ribosome profiling). We found evidence of translation for 40-50% of the expressed transcript isoforms in human and mouse, with 53% of the expressed genes having more than one translated isoform in human, 33% in mouse. Differential analysis revealed that about 40% of the splicing changes at RNA level were concordant with changes in translation, with 21.7% of changes at RNA level and 17.8% at translation level conserved between human and mouse. Furthermore, orthologous cassette exons preserving the directionality of the change were conserved between human and mouse and enriched in microexons in a comparison between glia and glioma. ORQAS leverages ribosome profiling to uncover a widespread and evolutionary conserved impact of differential splicing on the translation of isoforms and, in particular, of microexon-containing isoforms. ORQAS is available at https://github.com/comprna/orqas

## Introduction

The alternative processing of transcribed genomic loci through transcript initiation, splicing, and polyadenylation, determine the repertoire of RNA molecules in cells (Brown et al. 2014). Differential production of transcript isoforms, especially through the mechanism of alternative splicing, is crucial in multiple biological processes such as cell differentiation, acquisition of tissue-specific functions, and DNA repair (Fiszbein and Kornblihtt 2017; Baralle and Giudice 2017; Shkreta and Chabot 2015), as well as in multiple pathologies (Ward and Cooper 2010; Singh and Eyras 2017; Cummings et al. 2017). Although analysis of RNA sequencing (RNA-seq) data from multiple samples has indicated a large diversity of transcript molecules (Pertea et al. 2018), genes express mostly one single isoform in any given condition and this isoform may change across conditions (Gonzàlez-Porta et al. 2013; Sebestyén et al. 2015).

Computational and in-vitro studies have provided evidence that a change in relative isoform abundances can lead to the production of protein variants that impact the network of protein-protein interactions in different contexts (Buljan et al. 2012; Yang et al. 2016; Climente-González et al. 2017; Wojtowicz et al. 2007). In contrast, quantitative proteomics of naturally occurring proteins has identified much fewer protein variants than those predicted with RNA sequencing (Ezkurdia et al. 2015; Liu et al. 2017). Using state-of-the-art proteomics, it was recently shown that splicing changes at RNA level lead to changes in the sequence and abundance of proteins produced, although this was detected only for a limited number of transcripts (Liu et al. 2017). The difficulty in establishing a correspondence between transcript and protein variation may be due to limitations in current proteomics technologies, but also to the stability and translation regulation of transcripts (Maslon et al. 2014; Braun and Young 2014). Despite the evidence about its functional relevance (Baralle and Giudice 2017), it is still debated whether differential splicing leads to fundamentally different proteins and how widespread this might be (Tress et al. 2017a; Blencowe 2017; Tress et al. 2017b). Of particular interest are microexons, which can be as short as 3 nucleotides and carry out conserved neuronal-specific functions, and whose misregulation is linked to autism (Raj et al. 2014; Irimia et al. 2014; Quesnel-Vallieres et al. 2016). Despite their involvement in protein-protein interactions (Ellis et al. 2012; Irimia et al. 2014), the detection of protein variation associated to differential microexon inclusion using proteomics is currently challenging.

Sequencing of ribosome-protected RNA fragments, i.e. ribosome profiling, provides information on the messengers being translated in a cell. In particular, it allows the identification of multiple translated open reading frames (ORFs) in the same gene and the discovery of novel translated genes (Ingolia et al. 2009; Michel et al. 2012; Ruiz-Orera et al. 2014, 2015). However, ribosome profiling studies have been mainly oriented to gene-level analysis (Ingolia et al. 2009; Gonzalez et al. 2014; Ruiz-Orera et al. 2014). Recently, reads from ribosome profiling have been mapped across the exon-exon junctions of alternative splicing events (Weatheritt et al. 2016), suggesting that alternative splicing products may be engaged by ribosomes and potentially translated to produce different protein isoforms. A potential limitation of that approach is that ribosomal profiling reads also contain signals from native, non-ribosomal RNA-protein complexes (Ji et al. 2016), hence the mapping of ribosome reads to these regions may not necessarily be indicative of active translation. Additionally, ribosome activity is associated to signal periodicity and uniformity along open reading frames (Ji et al. 2015), which has not yet been tested in relation to transcript isoforms and alternative splicing. Thus, the extent to which alternative splicing, and in particular microexon inclusion, leads to the translation of alternative ORFs remains largely unknown.

In this article, we describe a new method, ORQAS (ORF quantification pipeline for alternative splicing), to quantify translation abundance at individual transcript level from ribosome profiling taking into account Ribosome signal periodicity and uniformity per isoform. We validate the translation quantification of isoforms using independent data from polysomal fractions and proteomics. We further find a concordance between differential splicing and differential translation and obtain evidence for the differential translation of microexons that is conserved between human and mouse. ORQAS provides a powerful strategy to study the impacts of differential RNA processing in translation.

## Results

### Translation Abundance estimation at isoform level from Ribo-seq

We developed a new method, ORQAS (ORF quantification pipeline for alternative splicing), for the estimation of isoform-specific translation abundance and to investigate the impact of differential splicing on translation (Fig. 1a) (Methods). ORQAS quantifies the abundance of open reading frames (ORFs) in RNA space from RNA sequencing (RNA-seq) in transcript per million (TPM) units, and assigns ribosome sequencing (Ribo-seq) reads to the same ORFs using RiboMap (Wang et al. 2016). After the assignment of Ribo-seq reads to isoform-specific ORFs, ORQAS only consider ORFs with at least 10 Ribo-seq reads after pooling replicates and with average RNA-seq abundance greater than 0.1 in transcript per million (TPM) units. ORQAS then calculates for each of these ORFs two essential metrics to determine their potential translation: Uniformity, calculated as a proportion of the maximum entropy of the read distribution, and the 3nt periodicity along the ORF. The translation abundance of each ORF is then calculated in ORF per million (OPM) units. These abundances are then used to study the impact of differential splicing on translation (Methods).

**Figure 1.**
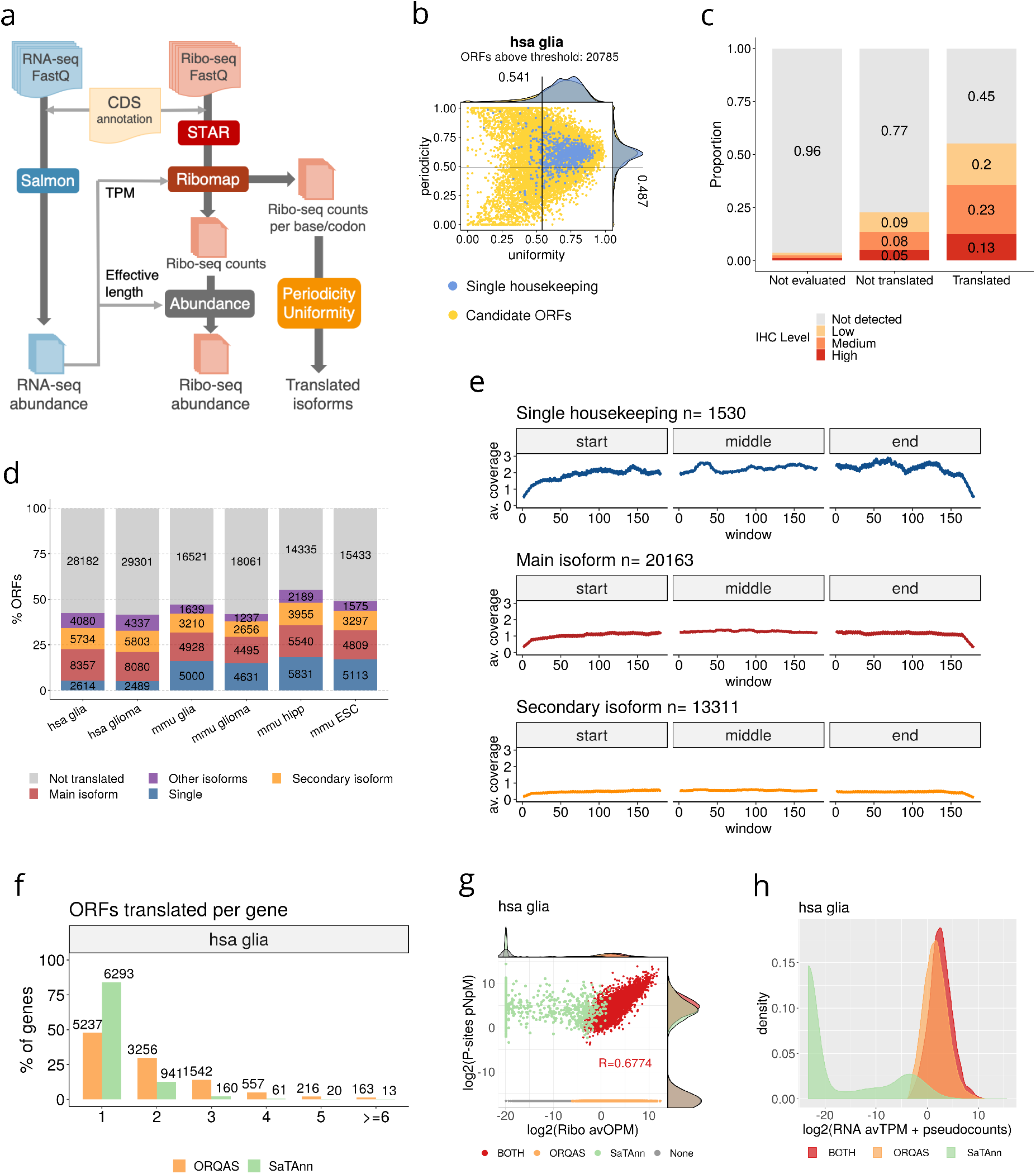
Estimation of translated isoforms. **(a)** The ORF quantification pipeline for alternative splicing (ORQAS) quantifies transcript abundances in transcripts per million (TPM) units from RNA-seq with Salmon (Patro et al. 2017) and in ORFs per million (OPM) units with Ribomap (Wang et al. 2016). Coverage and periodicity are calculated for every ORFs with TPM > 0.1, and candidate translated isoforms are estimated by comparing to a set of single-ORF house keeping genes with protein expression evidence across 37 tissues. **(b)** Uniformity (x axis) versus periodicity (y axis) for all tested ORFs with RNA expression TPM > 0.1 and at least 10 Ribo-seq reads assigned (in yellow). Uniformity is measured as the percentage of maximum entropy and periodicity is measured in the first annotated frame. Single-ORF genes with protein expression in 37 tissues from the Human Protein Atlas (THPA) are indicated in blue. We show the data for human glia (hsa glia). Other samples are shown in Supp. Fig. 1. **(c)** ORQAS prediction from human glia data on 1005 human genes annotated with one single ORF. The plot shows the level of protein expression from immunohistochemistry (IHC) experiments in human cortex from THPA for cases that do not show enough RNA expression to be evaluated by ORQAS (Not evaluated), cases that do not pass the filters of uniformity and periodicity (Not translated), and those predicted to be translated. **(d)** Number of ORFs predicted to be translated per sample, separated according to whether the ORF is encoded by: a single-ORF gene (Single), the most abundant isoform according to RNA-seq abundance in a gene with multiple isoforms (Main isoform), the second most abundant (Secondary isoform), or by any of the remaining isoforms in the abundance ranking (Other isoforms). Tested ORFs that are not predicted to be translated are depicted in gray (Not-translated). **(e)** Average density of Ribo-seq reads along ORFs in housekeeping singleton genes, in ORFs from the most abundant isoform according to RNA-seq abundance in a gene with multiple isoforms, and in the second most abundant isoform. **(f)** Distribution of the number of different ORFs translated per gene in the human glia (hsa glia) samples according to ORQAS (orange) and SaTAnn (green) predictions. Other samples are shown in Supp. Fig 6. **(g)** Correlation between the average ORF abundance in Ribosome space in the human glia (hsa glia) samples measured as ORFs per Million (OPM) in ORQAS and P-sites per Nucleotide per Million (P-sites pNpM) calculated from SaTAnn. Other samples are shown in Supp. Fig 8. Each ORF is plotted according to whether it is predicted only by ORQAS (orange), only by SaTAnn (green), or by both methods (red). **(h)** Distribution of the average RNA expression in the human glia (hsa glia) samples measured in TPM for ORFs that are predicted as translated only by ORQAS (orange), only by SaTAnn (green) or by both methods (red). Other samples are shown in (Supp. Fig 9).

We used ORQAS to analyze Ribo-seq and matched RNA-seq data from human and mouse glia and glioma (Gonzalez et al. 2014), mouse hippocampus (Cho et al. 2015), and mouse embryonic stem cells (Sugiyama et al. 2017) (Supp. Table 1). To determine which values of uniformity and periodicity are indicative of an isoform being translated, we selected as positive controls 929 human genes with a single annotated ORF and with evidence of protein expression from massspectrometry (MS), immunohistochemistry (IHC), and Uniprot in all 37 tissues available in the Human Protein Atlas (THPA) (Uhlén et al. 2015). For the mouse samples, the positive controls were 802 genes with one-to-one orthology with the human positive controls. We considered translated those ORFs within the 90% of the periodicity and uniformity distribution of these positive controls (Fig. 1b) (Supp. Fig. 1). This produced a total of 20709-20785 translated ORFs in human, and 13,019-17,515 in mouse (Supp. Table 2).

To determine the robustness of our filter on RNA abundance, we considered other cut-offs, but results did not change significantly after removing cases below 1 TPM (Supp. Fig. 2). As a further quality control, we considered the proportion of isoforms with low or no RNA expression that fell inside our periodicity and uniformity cutoffs and found only 0.7-0.9% across the human samples and 0.1-1.5% in the mouse samples (Supp. Table 3). To show that ORQAS provides an advantage over simply quantifying isoforms from Ribo-seq, we analyzed 1005 genes with one single annotated ORF annotated not included in the list of positive controls used above, and calculated the proportion of cases that showed evidence of protein expression from IHC experiments from THPA. We observed that cases that did not pass ORQAS thresholds generally lacked protein evidence (Fig. 1c). A similar analysis with all genome-wide predictions, not including any single-ORF gene, also showed that genes with translated isoforms are more frequently validated at all levels of protein expression, and that the majority (96%) of genes with translated ORFs showed some evidence of protein expression from either MS, IHC or Uniprot (Supp. Fig. 3a and 3b). To further validate the selection of these cut-offs, we considered genes with specific protein expression in brain, heart, intestine, liver, spleen or testis. We found that ORQAS predicts in glia a higher proportion of translated ORFs in the subset of brain-specific genes compared with the other tissues (Supp. Fig. 3c).

ORQAS predicted that a large fraction of the expressed protein-coding genes had multiple translated isoforms: 52,3%-54,9% of the genes in human and 29.1%-35.9% in mouse (Supp. Fig. 4). Overall, the majority of translated isoforms corresponded to either single-isoform genes or to the isoform with the highest expression in a sample (main isoform) (Fig. 1d). However, from those genes with multiple isoforms expressed at the RNA level, 3,471-3,570 (52.6%-55.5%) of genes in human and 577-898 (27.6-34%) in mouse had an alternative isoform translated (Fig. 1d). From all translated isoforms, 47.3%-49.2% in human, and 28.3%-34.9% in mouse, correspond to alternative isoforms (secondary or other isoform, Fig. 1d). In genes with multiple isoforms, the main isoform showed the highest average Ribo-seq coverage compared to secondary isoforms, albeit not as high as for the single-ORF genes used as positive controls (Fig. 1e). Additionally, the periodicity of the translated isoforms was uniform across the ORFs (Supp. Fig. 5).

We compared ORQAS with SaTAnn (Calviello et al. 2019), another method to calculate translation at isoform level from Ribo-seq. We run SaTAnn on the same samples analyzed with ORQAS. Looking at the annotated ORFs, ORQAS detected more genes with multiple ORFs consistently across all samples tested (Fig. 1f) (Supp. Fig. 6). Calculating the level of agreement between both methods with a Jaccard index, there was a higher level of agreement at gene level than at isoform level (Supp. Fig. 7). We also compared the translation abundance provided by both methods. SaTAnn abundance is based on the normalized number of P-sites per nucleotide, whereas ORQAS provides an abundance in ORFs per million (OPM), akin to TPM units. We observed in general a good correlation of the abundance values for ORFs predicted to be translated by both methods (Fig. 1g) (Supp. Fig. 8). However, many isoforms that did not pass ORQAS read-count filter still showed high abundance according to SaTAnn. ORFs predicted only by SaTAnn turned out to have low or no RNA-seq expression (Fig. 1h) (Supp. Fig. 9).

### Ribosome profiling discriminates translation abundance at isoform level

To validate the translation predictions by ORQAS at isoform level, we compared the results from Ribo-seq with the RNA abundances measured from polysomal fractions in the same human neuronal and embryonic stem cell samples (Blair et al. 2017). ORQAS predicted 27,552 translated isoforms in stem cells, and 25,034 in neurons (Supp. Fig. 10a). To control for the fact that monosomes can contain translating short mRNAs (Heyer and Moore 2016) (Supp. Fig. 10b and 10c), we separated isoforms in three different length ranges. We found that translated isoforms predicted by ORQAS were enriched in polysomal fractions at all length ranges (Fig. 2a) (Supp. Fig. 10d). In contrast, isoforms with RNA expression but not predicted to be translated with ORQAS were enriched in monosomal fractions (Fig. 2a). This provides support for our predictions and is also consistent with a small proportion of our predicted translated isoforms to be associated with non-sense mediated decay (NMD) targets, which are generally associated with monosomes (Kim et al. 2017).

**Figure 2.**
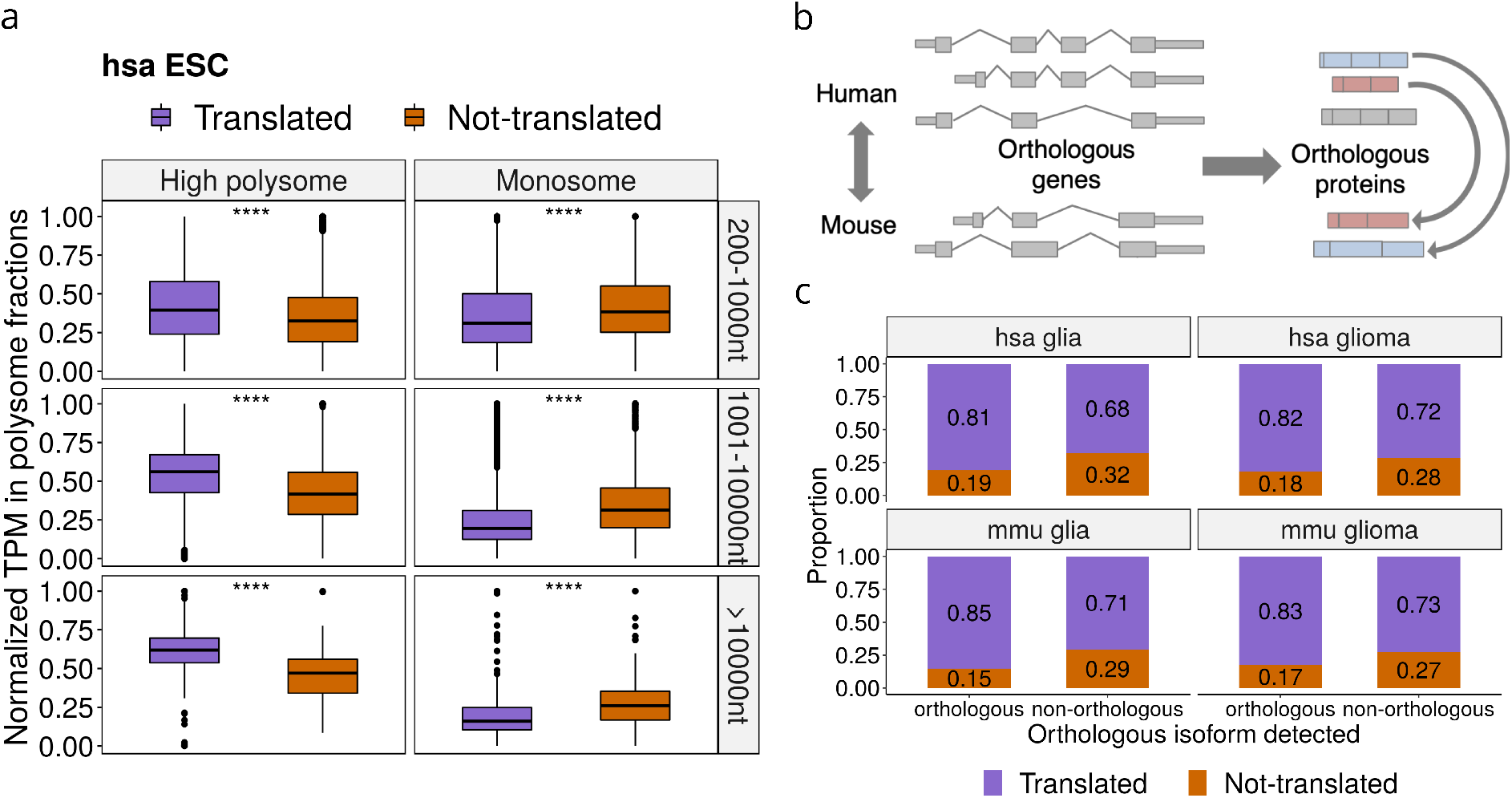
Validation of predictions. **(a)** We show the distribution of the relative abundance in high polysome (left panels) and monosome (right panels) fractions of human Embryonic Stem Cells (ESC) for translated isoforms and for isoforms with RNA expression (TPM>0.1) but predicted as not translated. The plot shows the results for three different ORF lengths: 200-1000nt (high polysome p-value 7.2e−67 and monosome p-value 2.3e−77), 1001-10000nt (high polysome p-value 2.5e−242 and monosome p-value 1.1e−94) and longer than 10000nt (high polysome p-value 5.1e−10 and monosome p-vaue 5e−06). The results for neuronal cells are given in Supp. Fig. 3. **(b)** Cross-species conservation of protein isoforms. Protein isoforms from a 1-to-1 orthologous gene pair are compared and candidate orthologous pairs are assigned using an optimization approach (Methods). **(c)** For the set of ORFs encoding a human-mouse orthologous protein pair (orthologous) and for those encoding proteins without an orthologous pair in mouse (non-orthologous) we plot the percentage that are predicted to be translated (translated) and the ones with RNA expression (TPM>0.1) but predicted as not translated (not-translated). We show here the results for human glia (p-value = 1.41e−140 Fisher test) and glioma (p-value = 3.63e−85), and for mouse glia (p-value = 1.143e−130) and glioma (p-value = 7.462e−53), Other mouse samples are shown in Supp. Fig. 10e.

Cross-species conservation is a strong indicator of stable protein production (Ezkurdia et al. 2014). We thus decided to test the conservation of our translated isoforms in human and mouse, using glia and glioma samples available for both species. To this end, we used an optimization method to determine the human-mouse protein isoform pairs most likely to be functional orthologs (Fig. 2b) (Methods). From 15,824 human-mouse 1-to-1 gene orthologs, we identified 18,574 human-mouse protein isoform pairs, and 7,112 (64%) of the 1-to-1 gene orthologs had more than one such isoform pair. These 18,574 protein pairs represent orthologous protein isoform pairs. We found that orthologous isoform pairs were significantly enriched in translated isoforms in both species (p-value < 2.2e−16 in all datasets) (Fig. 2c), providing further support for our predictions.

To perform an additional validation of our findings, we considered isoform-specific regions (Fig. 3a), since evidence mapped to these regions can then be unequivocally assigned to the isoform (see Supp. Figs. 11 and 12 for specific examples). We defined two types of isoform-specific regions. One type was defined in terms of isoform-specific nucleotide sequences, i.e. continuous nucleotide stretches that are only included in an isoform. From the annotation we were able to identify 3455 isoforms with such regions in human and 29447 in mouse. We found that translated isoforms had a significantly higher density of Ribo-seq reads per nucleotide in those regions than non-translated isoforms (Fig. 3b) (Supp. Fig. 13a). Additionally, unique sequence regions harbored more uniquely mapping Ribo-seq reads in translated isoforms compared to non-translated ones (Fig. 3c) (Supp Fig. 13b). Overall, we were able to validate 56-80% of the isoform-specific sequence regions.

**Figure 3.**
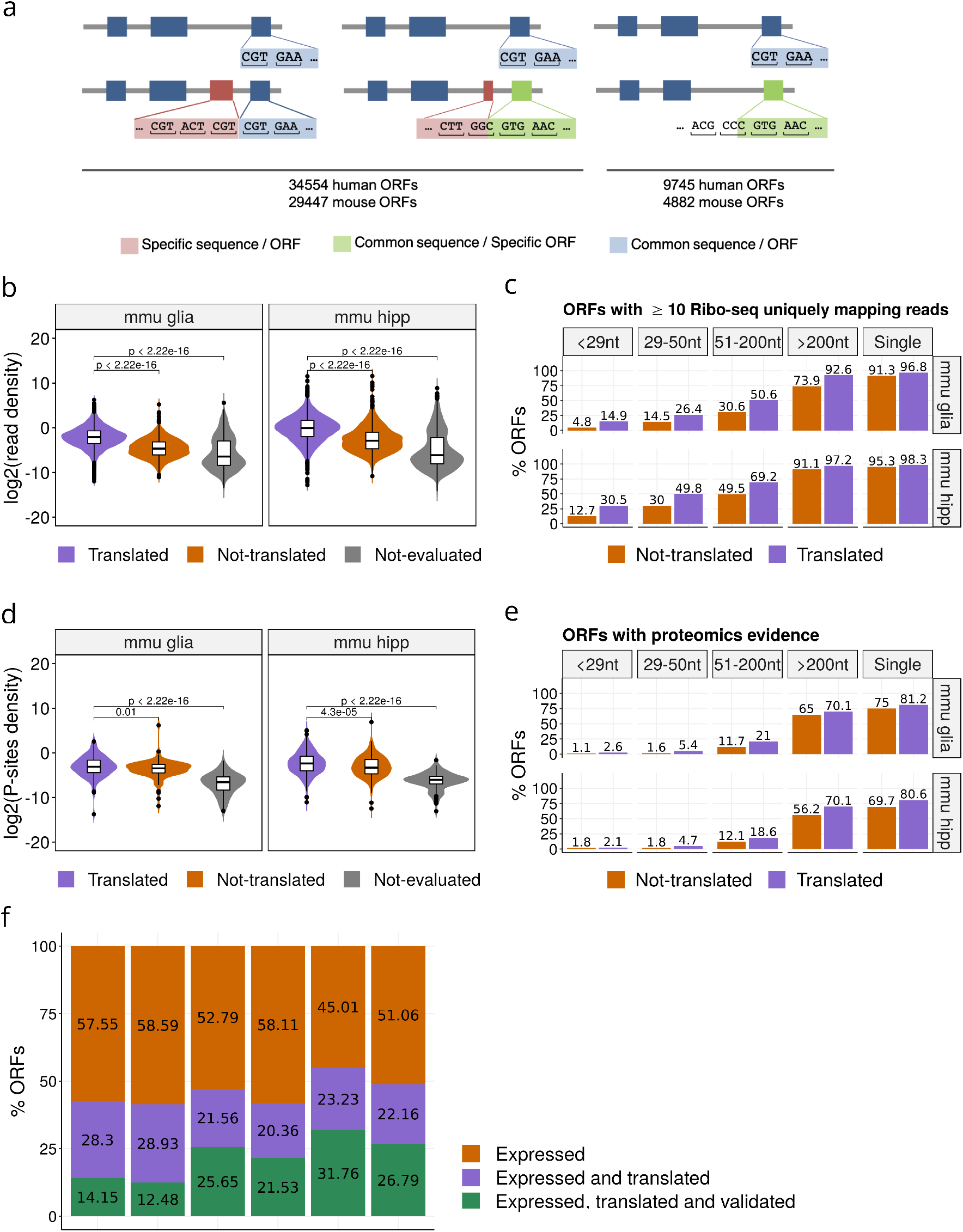
Validation with Isoform-specific regions. **(a)** Isoform-specific sequence regions (pink) are defined as the parts of an isoform ORF that are not present in any other isoform from the same gene. Isoform-specific ORFs (green) are defined as a region shared between two isoforms, but with a different frame in each isoform. **(b)** For the mouse samples of glia (mmu glia) and hippocampus (mmu hipp) we show the density of Ribo-seq reads per nucleotide over the isoform-specific sequence regions, calculated as the uniquely mapping read-count over region length in log_2_ scale for isoform-specific sequences. Distributions are given for predicted translated isoforms, for isoforms that did not pass the threshold of uniformity and periodicity (not translated), and for the isoforms with low expression (TPM<0.1) (not evaluated). Other samples are shown in (Supp Fig. 13a). **(c)** For the mouse samples of glia (mmu glia) and hippocampus (mmu hipp) the plot shows the percentage of regions with at least 10 uniquely mapping Ribo-seq reads in isoform-specific sequences over the total number of isoforms with an isoform-specific sequence defined according to the length of the region. Other samples are shown in (Supp Fig. 13b). **(d)** For the mouse samples of glia (mmu glia) and hippocampus (mmu hipp) we show the density of Ribo-seq reads per nucleotide over the isoform-specific ORFs, calculated as the counts per nucleotide based on the estimated P-site positions over region length in log_2_ scale for isoform-specific ORFs. Other samples are shown in (Supp Fig. 13c). **(e)** For the mouse samples of glia (mmu glia) and hippocampus (mmu hipp), the plot shows the percentage of ORF-specific regions with 1 or more mass-spectrometry peptides, separated according to region length. The plot shows the combined results for both types of regions: isoform-specific sequence regions and isoform-specific ORFs. **(f)** Proportion of isoforms expressed predicted to be translated and that have been validated using one or more sources of evidence: conservation, uniquely mapped Ribo-seq reads in isoform-specific sequences, and counts per base or peptides in isoform-specific ORFs. For each sample, and for all ORFs with sufficient RNA-seq expression (TPM > 0.1), we show the proportion predicted to be translated from Ribo-seq reads and the proportion that were validated. The proportions are colored to indicate the fraction that is not included in the more restricted set: expressed > translated > validated.

To be able to validate our predictions using P-sites and peptides from MS experiments, we additionally considered isoform-specific ORF regions (Fig. 3a). These were defined as sequences that may or may not be shared between isoforms but had a specific frame in each isoform, so that peptides from MS experiments can be unequivocally mapped on these regions. From the annotation we found in total 44,299 isoforms with specific ORF regions in human and 34,329 in mouse. These included the 34,554 and 29,447 isoforms calculated before with ORFs from isoform-specific sequences in human and mouse, respectively: hence, there were 9745 and 4882 isoforms in the human and mouse annotations, respectively, that did not have differences in sequence but had different overlapping ORFs. We found that translated isoforms had a significantly higher density of P-sites in isoform-specific ORFs (Fig. 3d) (Supp. Fig. 13c). We further used peptides from MS experiments ^42^ to validate our predictions. Overall we validated more isoforms predicted as translated compared with those predicted as not translated (Fig 3e). The rate of validation decreased with the region length, as expected for MS experiments ^41^. On the other hand, the proportion of predictions validated with peptides increased using increasing cut-offs for RNA expression (Supp. Fig. 13d). Overall, we were able to validate 48%-73% of the isoform-specific ORFs tested.

In summary, from all the protein-coding transcript isoforms considered from the annotation (84,024 in human and 48,928 in mouse), 58-59% in human and 63-65% in mouse showed RNA expression > 0.1 TPM (Supp. Table 4) (translated isoforms are available in Supplementary Data File 1). From these expressed isoforms, about 40% in human and 41-54% in mouse were predicted to be translated by ORQAS, and 23-43% were validated using independent data, including conservation (Fig. 3f). Furthermore, 60% of the alternative isoforms predicted as translated had independent evidence of translation, and these corresponded to approximately 10% of all the annotated alternative isoforms in human and mouse (Supp. Fig. 13e). Our analyses thus indicate that alternative transcript isoforms are often translated into protein, although they represent a small fraction of all expressed transcripts.

### Conserved impact of differential splicing on translation

Differential splicing is often assumed to lead to a measurable difference in protein production. However, this has only been shown for a limited number of cases (Liu et al. 2017). We addressed this question at genome scale using our more sensitive approach based on Ribo-seq. We used SUPPA (Alamancos et al. 2015; Trincado et al. 2018) to obtain 37,676 alternative splicing events in human and 17,339 in mouse that covered protein coding regions (Methods) (all alternative splicing events are available in Supplementary Data File 2). Using the same SUPPA engine to convert isoform abundances to event inclusion values (Alamancos et al. 2015; Trincado et al. 2018), we calculated a relative abundance (RA), defined as the proportion of translation abundance of the isoforms given by ORQAS that is explained by a particular alternative splicing event (Fig. 4a). Accordingly, in analogy to a relative inclusion change (ΔPSI) in RNA space, we were able to measure the relative differences in ribosome space due to the inclusion or exclusion of particular alternative exons, or ΔRA.

**Figure 4.**
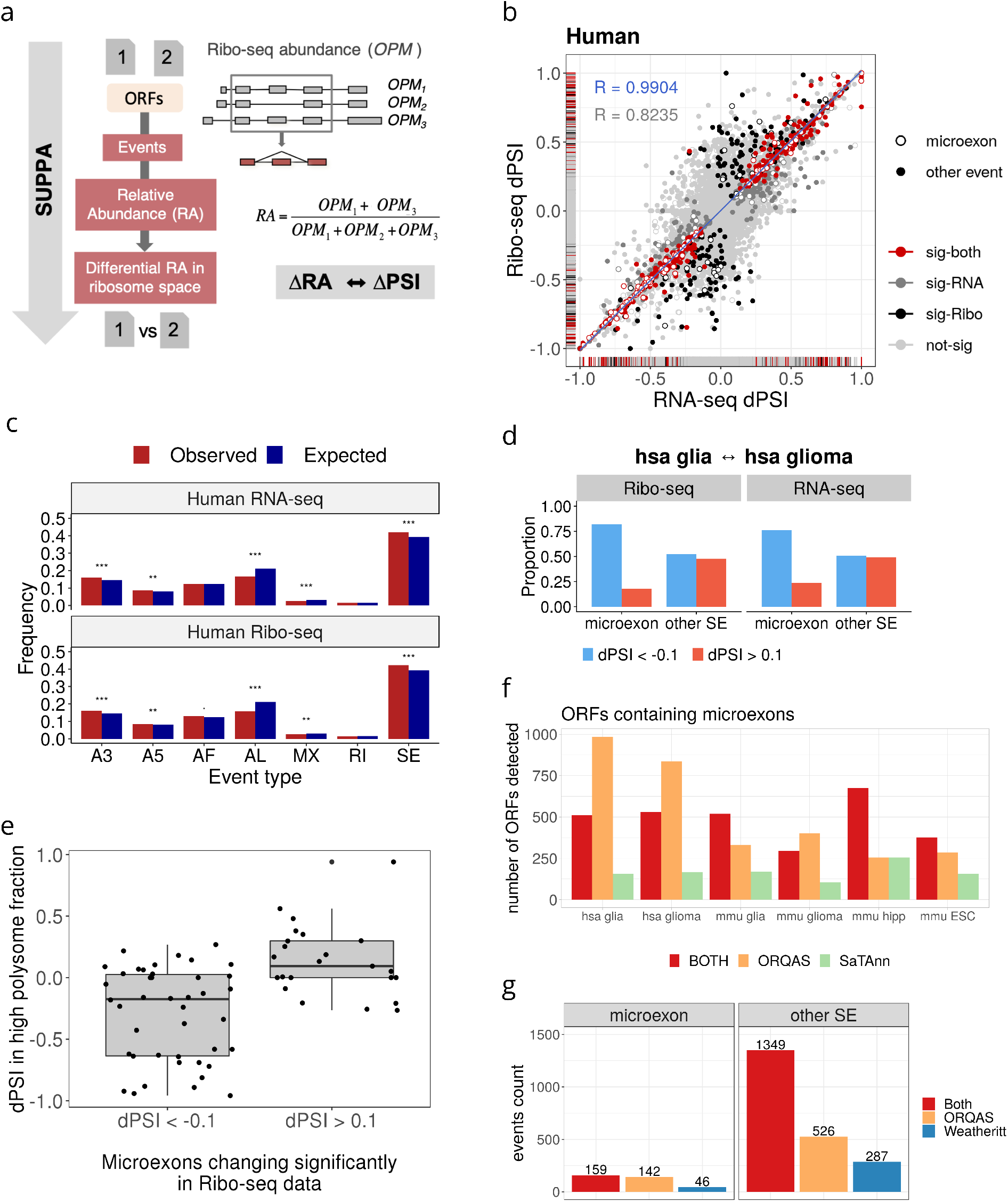
Differential translation linked to differential splicing. **(a)** Description of how SUPPA was used to calculate the differential inclusion of events in Ribosome space. The abundance of open reading frames (ORFs) is calculated in ORFs per million (OPM) units. OPMs are transferred to event inclusion values using SUPPA definition of events, using only exon-intron structures overlapping ORFs. For each event a relative abundance (RA) is obtained, analogous to a PSI. **(b)** Correlation of changes in splicing and translation in events in human. The figure shows in red the cases that are significant in both Ribosome and RNA space (363 events), in black (493?) or dark gray (227), the cases that are only significant in one case, and in light gray, the cases that are significant in neither of them. The density bands in the plot show the distribution of cases along the axes. Empty circles indicate the microexons (red if significant, black/dark-gray/gray otherwise). In the inset in blue we give the correlation of the red points (including microexons), in gray we give the correlation of all other exons (black/dark-gray/gray, including microexons). **(c)** We show the differentially spliced events calculated in RNA space (upper panel) or in Ribosome space (lower panel) and separated by event type. In blue we show the proportion of events calculated with SUPPA from the annotated coding regions in transcripts in human, whereas in red we show the proportion of events with a significant change between human glia and human glioma. Event types are: alternative 3’ss (A3) and 5’ss (A5), alternative first (AF) and last (AL) exon, mutually exclusive (MX) exon, retained introns (RI) and skipping exon (SE). There is significant enrichment for SE events for RNA (Fisher’s test p-value = 3.90e−06) and Ribo-seq (p-value = 7.46e−07), for A3 events for RNA (p-value = 1.02e−05) and Ribo-seq (6.22e−06), for A5 events for RNA (4.55e−03) and Ribo-seq (8.73e−03); and significant depletion for AL events for RNA (1.73e−32) and Ribo-seq (2.43e−43) and for MX events for RNA (6.06e−04) and Ribo-seq (1.83e−03) **(d)** Enrichment of microexons with an impact in RNA splicing and ORF translation in human from the comparison of glia and glioma samples. In the figure, dPSI is used to indicate the difference in relative abundance in both RNA and Ribosome spaces. **(e)** Difference in high polysome fraction, measured as dPSI, between neuronal samples and ESCs (y axis) for microexons with a significant change in Ribosome space. As before, dPSI indicates the difference in relative abundance in Ribosome space. **(f)** Number of ORFs containing microexons that are detected only by ORQAS (orange), only by SaTAnn (green) or by both methods (red). **(g)** Number of microexons (<52nt) and longer SE exon events (other SE) that are detected in Ribosome space with PSI > 0.1 depending on the method: applying ORQAS pipeline followed by SUPPA (orange), using the results from (Weatheritt et al. 2016) obtained by mapping reads to exon-exon junctions (blue), or by both methods (red).

Comparing the glia and glioma samples in human, we found 856 events with a significant change in RNA splicing (|ΔPSI|>0.1 and p-value<0.05), and 590 events with significant differential translation (|ΔRA|>0.1 and p-value<0.05), with a significant overlap of 363 events between them (Jaccard index, z-score=89.386 comparing with the Jaccard index distribution of the overlaps from 1000 subsample sets of the same size) (Supp. Fig. 14a). Similarly, in mouse we found an overlap of 179 events (Jaccard index z-score=65.326) between 471 events with a significant change in RNA splicing (|ΔPSI|>0.1 and p-value<0.05) and 240 with significant change in translation (|ΔRA|>0.1 and p-value<0.05) (Supp. Fig. 14b).

We observed a concordance in the direction and magnitude of the change in significant events in RNA and ribosome space in human (Pearson R=0.9904, p-value = 5.309e−312) and mouse (Pearson R=0.9937, p-value = 2.113e−170) and this concordance was greater than for the rest of events (Fig. 4b) (Supp. Fig. 14c). We further observed a similar proportion of event types changing significantly in RNA and ribosome space, with an enrichment of exon skipping events in human (Fig. 4c) and mouse (Supp. Fig. 14d). To investigate the nature of these enriched cases, we considered very short alternative exons, or microexons, which are known to be differentially included in brain cells (Irimia et al. 2014; Li et al. 2015). We tested the inclusion properties of alternative exons in RNA and Ribosome space in our samples. We observed that alternative exons show less inclusion in glioma compared with glia at lengths 51nt or below, with a stronger pattern below 28nt, which are the two previously employed length cut-offs to define microexons (Supp. Fig. 14e) (Irimia et al. 2014; Li et al. 2015).

Microexons, defined as exons of length 51nt or less, were enriched in the events with significant changes in RNA-seq (Fisher’s test p-value 0.01125200 in human, 1.231482e−10 in mouse) and in Ribo-seq (Fisher’s test p-values 0.03917744 in human, 1.049473e−06 in Mouse). This enrichment was also present if microexons were defined to be of length <28nt: Glia vs Glioma in human (Fisher’s test p-value for RNA-seq 5.435e−13, for Ribo-seq 1.17e−09), Glia vs Glioma in mouse (Fisher’s test p-value for RNA-seq 7.47e−14, for Ribo-seq 3.194e−06), Human ESC vs differentiated neurons (Fisher’s test p-value for RNA-seq 2.725e−08, for Ribo-seq 6.768e−06).

Moreover, microexons were enriched in events decreasing inclusion in glioma compared with glia in both human (Fisher’s test p-values 1.382e−12 for RNA-seq and 5.602e−10 for Ribo-seq) (Fig. 4d) and mouse (Fisher’s test p-values 6.386e−16 for RNA-seq and 3.446e−06 for Ribo-seq) (Supp. Fig. 14f). We repeated the same analysis using data from human neuronal differentiation (Blair et al. 2017) and found that microexons were also enriched in the comparison between embryonic stem cells and neuronal cells in terms of RNA splicing and translation (p-values 8.435e−06 for RNA-seq and 6.597e−05 for Ribo-seq) (Supp. Fig. 14g). Furthermore, using RNA sequencing from polysome fractions from the same stem cell and neuronal samples, we were able to validate the change in inclusion patterns of microexons under the same conditions (Fig. 4e).

We compared the capacity of ORQAS and SaTAnn to detect translation of microexon-containing ORFs. In general, we observed that isoforms predicted by ORQAS included more cases with short (<51nt) and very short (<29nt) isoform-specific regions than SaTAnn (Supp. Fig. 15), hence providing a greater potential to identify translating microexons. Indeed, ORQAS identified more microexons than SaTAnn in all tested samples (Fig. 4f). We also performed a comparison with previously detected alternative splicing events using a direct mapping of Ribo-seq reads to the exon-exon junctions (Weatheritt et al. 2016). While there was a high overlap for exons longer than 51nt, ORQAS identified more microexons (Fig. 4g).

Our results provide evidence that differential splicing leads to a qualitative and quantitative change in the protein products from a gene locus. These results are also consistent with a functional relevance of the inclusion of microexons in protein-coding transcripts in neuronal differentiation and their inclusion loss in brain-related disorders (Raj et al. 2014; Irimia et al. 2014). Our analyses also highlight the strength of ORQAS in detecting these microexons compared with other alternative methods.

To further test the relevance of our findings, we considered a set of 1,487 alternative exons conserved between human and mouse (Fig. 5a). A high proportion of them changed in the same direction between glia and glioma (66% in RNA-seq and 78% in Ribo-seq). Moreover, we observed that among the events with concordant changes between both species there was an enrichment of microexons, with a significant trend towards less inclusion in glioma (p-value 5.389e−05 for RNA-seq and 5.521e−04 for Ribo-seq) (Fig. 5b). Furthermore, there was a correlation between the changes in Ribosome space in human and mouse (Fig. 5c).

**Figure 5.**
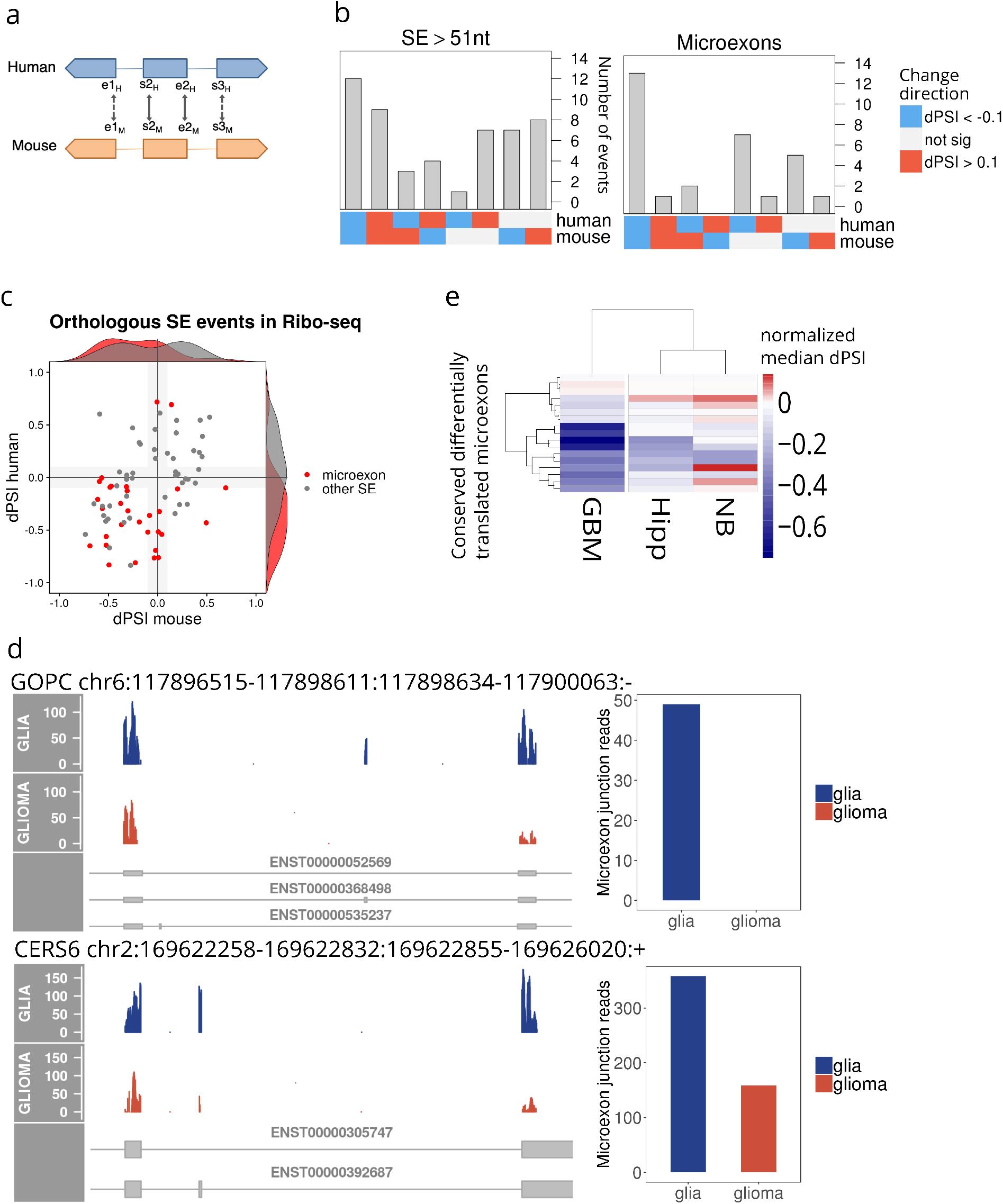
A conserved program of differential RNA splicing and translation. **(a)** Conserved alternative splicing events were obtained by mapping with LiftOver the coordinates of the alternative exon (s2, e2), and the internal coordinates of the flanking exons (e1, s3). We considered conserved those alternative exons that had at least (s2, e2) conserved. **(b)** Directionality of the changes in conserved alternative exons longer than 51nt (left panel) and microexons (right panel) in Ribosome space. As before, dPSI indicates here the difference in relative abundance in both RNA and Ribosome space. The plot shows the number of events (y axis) according to whether they were significant in human, mouse, or both, and the direction of change (x axis). We indicate in blue if the event had a significant decrease in inclusion smaller than −0.1 and in read if the event had a significant increase in inclusion larger than 0.1. **(c)** Correlation between the difference in relative abundance (dPSI) in human on the y axis and mouse on the x axis for the conserved differentially spiced events in Ribosome space. Microexons are depicted in red. **(d)** Examples of microexons that change significantly in RNA and ribosome space between glia and glioma. For the genes *GOPC* and *CERS6* we show Ribo-seq reads mapping to the microexon region and its flanking exons (left panels) and the number of Ribo-seq reads crossing the microexon junctions (right panels) in both glia (in blue) and glioma (in orange). **(e)** Patterns of inclusion of conserved differentially translated microexons in normal hippocampus (Hipp) samples from GTEX, gliblastoma multiforme (GBM) from TCGA, and neuroblastoma (NB) from TARGET. The heatmap shows the difference of the median PSI with respect to normal brain cortex tissue from GTEX.

Among the microexons with a differential pattern of splicing and translation, we identified one in the gene *GOPC* (Fig. 5d), which was previously linked to glioblastoma (Charest et al. 2003), and one in the gene *CERS6* (Fig. 5d), which has been associated with chemotherapy resistance (Verlekar et al. 2018). To test further the potential relevance of the identified microexons with conserved differential pattern, we calculated their RNA splicing inclusion patterns across other normal and tumor brain samples. In particular, we analyzed samples from glioblastoma multiforme (GBM) from TCGA (Cancer Genome Atlas Research Network 2008), neuroblastoma (NB) from TARGET (Pugh et al. 2013) (Fig 5e), and samples from cortex and hippocampus from GTEX (Carithers et al. 2015). Microexons with a conserved impact on translation recapitulate the pattern of decreased inclusion in GBM compared with the postmortem normal brain regions (Fig. 5e). Differentially translated microexons may explain tissue differentiation as well as tumor specific properties, as they separate out healthy tissues and tumor types (Supp. Fig. 16a). In contrast, conserved microexons are clearly more prevalent in differentiated brain tissues than in tumor samples (Supp. Fig. 16b).

## Discussion

We have described a new method, ORQAS (https://github.com/comprna/orqas), to obtain transcript abundance estimates at isoform level in ribosome space, to identify multiple protein products from a gene, and to investigate differential translation associated with alternative splicing and differential transcript usage between conditions. Our approach presents several novelties and advantages over other methods (Weatheritt et al. 2016; Calviello et al. 2019; Wang et al. 2016; Blair et al. 2017): *i)* filtering based on the periodicity and uniformity of the Ribo-seq reads improves the identification of translated isoforms; *ii)* the use of RNA expression to guide the translation abundance estimation avoids many potential false positives; *iii)* our description in terms of isoforms allows the identification of more translated alternative isoforms and, in particular, microexons compared with other methods, *iv)* our validation using isoform-specific regions, regardless of whether these regions could be encoded into an standard alternative splicing event, provides a more general validation of the impact of differential splicing on translation; and *v)* ORQAS provides an isoform abundance estimates in ribosome space that can be reused by other tools, like SUPPA.

We estimated that in total about 40-50% of the protein coding isoforms with RNA expression showed some evidence of translation, and that around 20,700 proteins are produced in human and 13,000-17,500 in mouse in the tested conditions. Additionally, about 5,700-5,800 genes in human and 2,600-3,900 in mouse produce more than one protein in those conditions. These estimates are considerably lower than what is generally predicted from RNA expression (Pertea et al. 2018). This may be explained by the limited coverage of Ribo-seq reads but may be also due to the fact that RNA-seq artificially amplifies fragments of unproductive RNAs leading to many false positives. Nonetheless, our data indicates that many more ORFs are translated in a given sample than what is detectable by current proteomics methods and that the number of detected translated ORFs is close to estimates obtained using a combination of proteomics and sequence conservation (Ezkurdia et al. 2014). Importantly, we found that multiple ORFs are translated from the same gene and at different abundances across conditions.

Around 40% of the events detected with differential RNA splicing showed consistent measurable changes in Ribo-seq in the same direction, which supports the notion that changes in RNA processing have a widespread impact in the translation of ORFs from a gene. In particular, we found that a pattern of decreased inclusion of microexons in glioma with respect to normal brain samples is recapitulated at the translation level, providing *in vivo* evidence that the splicing changes in microexons have an impact in protein production. Microexon inclusion is a hallmark of neuronal differentiation (Irimia et al. 2014; Raj et al. 2014; Trincado et al. 2018), and glia partly recapitulates the pattern of microexon inclusion found in neurons (Irimia et al. 2014). The decreased inclusion of microexons observed in glioma suggests a dedifferentiation pattern similar to the one previously described before for other tumors (Sebestyén et al. 2016), but could also be indicative of a difference in the content of neuronal cells in the samples compared. In either case, the evolutionary conservation of the change at RNA expression and protein production indicates a conserved functional program in these samples.

Our capacity to predict RNA splicing variations from RNA-seq data currently exceeds our power to evaluate the significance of those events regarding protein production with current proteomics technologies (Wang et al. 2018). Despite this limitation, mass spectrometry can show for a small number of cases that splicing changes impact the abundance of proteins produced by a gene (Liu et al. 2017). Our findings are in agreement with these results, and moreover overcome current limitations to determine genome-wide impacts of RNA processing changes on protein production. Furthermore, our analyses indicate that the majority of translated alternative isoforms shows less than 25% variation in length with respect to the most highly expressed isoform, suggesting that for most part, the functional impacts from alternative splicing are mediated by slight modifications in the protein sequences (Ellis et al. 2012), rather than through the production of essentially different proteins. In summary, ORQAS leverages ribosome profiling to provide a genome-wide coverage of genes and transcript isoforms and allow a more effective testing of the impacts of splicing in protein production, as well as the identification and validation of multiple proteins from the same gene locus.

## Methods

### Pre-processing of RNA-seq and Ribo-seq datasets

RNA-seq and Ribo-seq datasets were downloaded from Gene Expression Omnibus (GEO) for the following samples: normal glia and glioma from human and mouse (GSE51424) (Gonzalez et al. 2014), mouse hippocampus (GSE72064) (Cho et al. 2015), mouse embryonic stem cells (GSE89011) (Sugiyama et al. 2017), and three steps of forebrain neuronal differentiation in human (GSE100007) (Blair et al. 2017). Adapters in RNA-seq and Ribo-seq datasets were trimmed using cutadapt v.1.12 with additional quality filters for RNA-seq (−q = 30). We further discarded reads that mapped to annotated rRNAs and tRNAs. Remaining reads in RNA-seq and Ribo-seq datasets were filtered by length (>= 26 nucleotides).

### Quantification of transcripts and open reading frames

We used the Ensembl annotation v85 for human (hg19) (GRCh37.85) and mouse (mm10) (GRCm38.85) removing pseudogenes, short isoforms (< 200 nt) and annotated isoforms with incomplete 5’ or 3’ regions. For the analysis of RNA-seq data we used Salmon v0.7.2 (Patro et al. 2017) to quantify transcript abundances in transcripts per million (TPM) units using the annotation of unique open reading frames (ORFs). To quantify coding sequences (CDS) at the isoform level with the Ribo-seq data we applied a modified version of Ribomap (Wang et al. 2016) that uses the RNA-seq abundance as priors for the optimization algorithm to distribute Ribo-seq reads among the different isoforms. As default, Ribomap uses the RNA-seq reads aligned to the transcriptome sequences with STAR (Dobin et al. 2013). Instead, we provided a direct quantification of the ORFs with RNA-seq using Salmon, to be used as priors by RiboMap. We calculated the translation abundances of each ORF based on Ribo-seq reads in ORFs Per Million (OPM) units, similar to TPM units, but for ORFs instead of complete transcripts and using Ribo-seq reads. OPM values are calculated as follows:

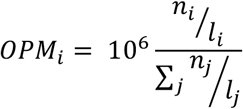

where *n*_*i*_ is the estimate Ribo-seq counts in ORF *i* and *l*_*i*_ is the effective length of the same ORF.

### Identification of actively translated isoform coding sequences

We identified actively translated ORFs by calculating two parameters read periodicity and read uniformity (Ji et al. 2015). The periodicity is based on the distribution of the reads in the annotated frame and the two alternative ones. In order to calculate the read periodicity, we previously computed the position of the P-site, corresponding to the tRNA binding-site in the ribosome complex. This was obtained by calculating the distance between annotated ATG start codons and the leftmost position covered by Ribo-Seq reads, for each read length. The uniformity was measured as the proportion of maximum entropy (PME) defined by the distribution of reads along the ORF:

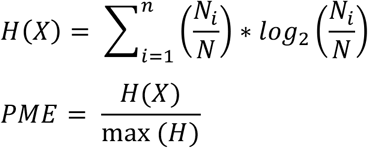

Where *N* represents the total number of reads, *N*_*i* i_is the number of reads in each region *i* and *max(H)* is the entropy assuming that the reads are equally distributed across the ORF. The maximum value is 1 and indicates a completely even distribution of reads across codons. These values were obtained for each sample by pooling the replicates and only considering ORFs with 10 or more assigned Ribo-seq reads, and with RNA-seq abundance TPM > 0.1.

### Polysomal fraction analysis

We used RNA-seq from high polysomal, low polysomal and monosomal fractions from embryonic stem cells and neuronal cell culture in human (GSE100007) (Blair et al. 2017) to quantify isoforms with Salmon (Patro et al. 2017). Only ORFs from protein-coding isoforms were used for quantification. For each isoform we calculated the polysomal relative abundance as before (Maslon et al. 2014) by dividing the abundance in each polysomal fraction in TPM units, by the sum of abundances in (high and low) polysomes and monosomes.

### Validation of isoform-specific regions

We defined two different types of isoform-specific regions that were analyzed differently. Isoform-specific sequences are regions with a unique nucleotide sequence among the isoforms of the same gene. Isoform-specific ORFs are defined as regions that will give rise to different amino-acid sequences within the proteins of the same gene, either because of the presence of isoform-specific sequences or frame-shifted common sequences (Fig. 3a). According to the annotation, we identified 34553 isoforms containing isoform-specific sequences in human and 29447 in mouse and 44298 isoforms containing isoform-specific ORFs in human and 34329 in mouse. For the validation of isoform-specific sequences we considered uniquely mapping Ribo-seq reads from the STAR output falling entirely inside these regions or in the junction of the specific sequence with the common region. Read densities inside those regions where calculated as the total number of uniquely mapping reads in the region divided by the length of the isoform-specific sequence. The validation of isoform-specific ORFs instead was performed using the profiles of counts in each base of the ORF considering the expected position of the P-site. For isoform-specific ORFs the read densities where established as total number of counts in the region divided by the length in nucleotides of the isoform-specific ORFs.

### Proteomics evidence in translated isoform coding sequences

We mined the proteomics database PRIDE (Vizcaino et al. 2016) to search for peptide matches to ORFs. We only considered peptide datasets from mouse corresponding to tissues analyzed in this study: brain (PRD000010, PXD000349, PXD001786), hippocampus (PRD000363, PXD000311, PXD001135), and embryonic cell lines (PRD000522). This corresponded to a total of 328,200 peptides. We searched for peptide matches in translated ORFs and only kept peptides that had one perfect match to an ORF and did not have a match with 0, 1 or 2 amino acid mismatches to any other annotated ORF isoform from the same or different genes.

### Differential inclusion of events at RNA and translation level

We used SUPPA (Alamancos et al. 2015; Trincado et al. 2018) to generate alternative splicing events defined from protein-coding transcripts and covering the annotated ORFs. The relative inclusion of an event was calculated as a Percent Spliced In (PSI) value with SUPPA in terms of the transcript abundances (in TPM units) calculated from RNA-seq and in terms of the ORF abundances (in OPM units) calculated from Ribo-seq (Fig. 4A). The test for significant differential inclusion of the events was applied in the same way for both cases by testing the difference between the observed change between conditions and the observed change between replicates, as described before (Trincado et al. 2018).

### Calculation of orthologous isoforms

We considered the set of 1-to-1 orthologous genes between human and mouse from Ensembl (v85) (Clamp et al. 2003). For each pair of orthologous genes, we calculated all possible pairwise global alignments between the human and mouse proteins encoded by these genes using MUSCLE (Edgar 2004). For each alignment we defined a score as the fraction of amino acid matches over the total length of the global alignment and kept only protein pairs with score >= 0.8. From all the remaining protein pairs in each orthologous gene pair, we selected the best human-mouse protein pairs using a symmetric version of the stable marriage algorithm (Eyras et al. 2004).

### Comparison with normal and tumor tissues

We downloaded the transcriptome (GRCh38) TPM values calculated with RSEM (Li and Dewey 2011) from the XENA browser (https://xenabrowser.net/datapages/) for GTEX (Carithers et al. 2015), TARGET (Pugh et al. 2013), and TCGA (Cancer Genome Atlas Research Network 2008). PSI values for the alternative splicing events in human were calculated with SUPPA (Alamancos et al. 2015; Trincado et al. 2018) from these TPM values, and the coordinates of the events were transformed to GRCh37 (hg19). We used 105 cortex and 84 hippocampus samples from GTEX, 171 Glioblastoma Multiforme samples from TCGA, and 162 Neuroblastoma samples from TCGA to analyze the PSI values of the identified translated microexons.

## Data availability

Predicted translated isoforms in the samples analyzed together with their validation using independent datasets are provided in Supplementary Data File 1. All calculated alternative splicing events in RNA and Ribosome space with annotations for microexons and orthology are provided in Supplementary Data File 2.

## Acknowledgements

We are grateful to T. Preiss and N. Shirokikh for useful discussions, to R. Weatheritt for providing access to his datasets, and to A. Closa for help obtaining the event data from cancer and normal tissue samples. We acknowledge funding from the Spanish Government and FEDER with grants BFU2015-65235-P, BIO2017-85364-R and MDM-2014-0370, and by Catalan Government (AGAUR) with grant SGR2017-1020. MR-S had funding from an FI grant from the Catalan Government with reference 2018FI_B1_00126 for part of this work.

