## Supplementary Figures for "Ribosome profiling at isoform level reveals an evolutionary conserved impact of differential splicing on the proteome"

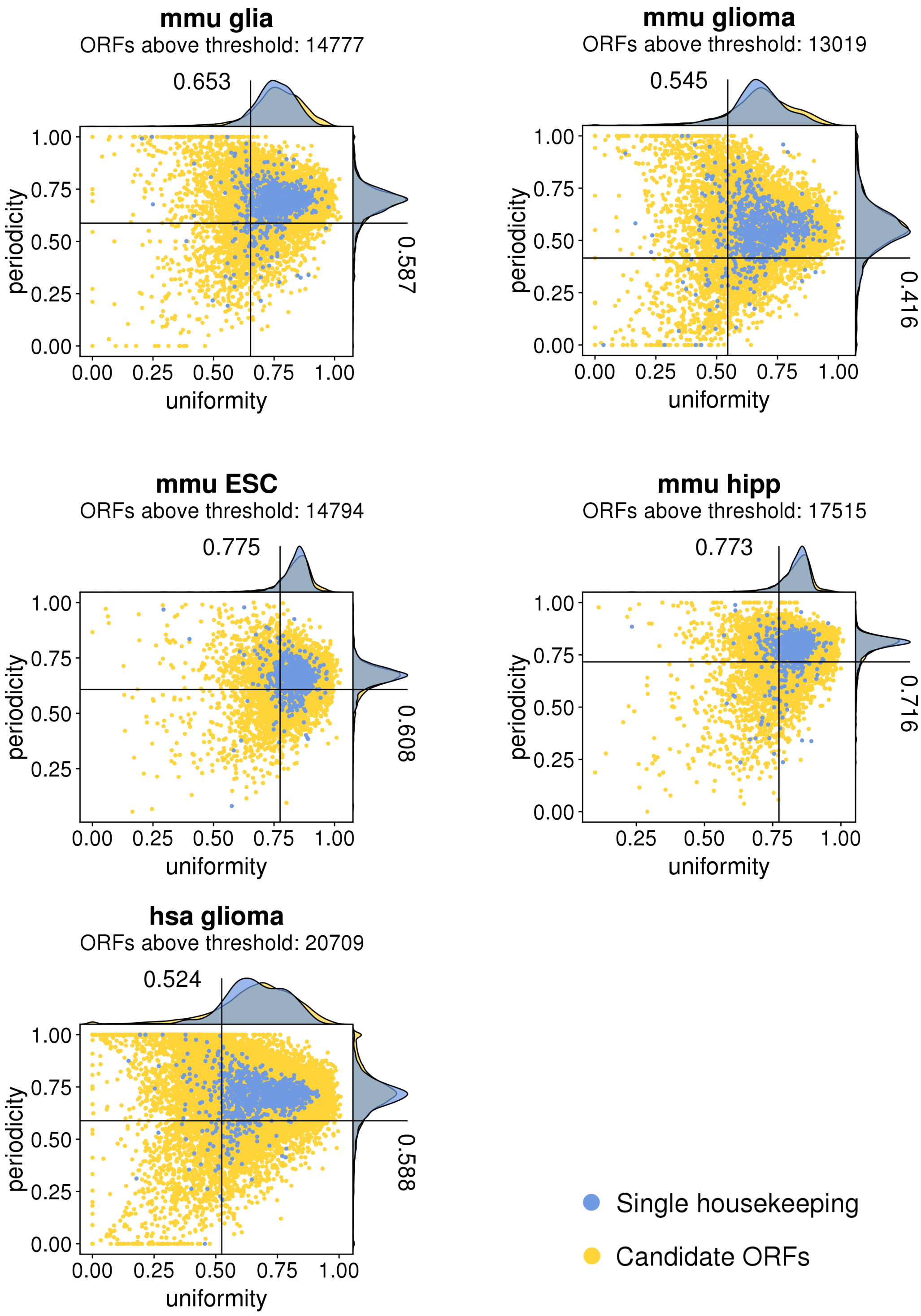


**Supplementary Figure 1.** Uniformity (x axis) versus periodicity (y axis) for ORFs with RNA expression TPM > 0.1 and at least 10 Ribo-seq reads. In blue we indicate ORFs from single-ORF genes with protein expression in all 37 tissues from THPA, and in yellow the rest of ORFs. Uniformity is measured as the percentage of maximum entropy and periodicity is measured in the first annotated frame. We show the data for the glioma human sample (hsa glioma) and the mouse samples from glia (mmu glia), glioma (mmu glioma), Embrionic Stem Cells (mmu ESC) and hippocampus (mmu hipp).

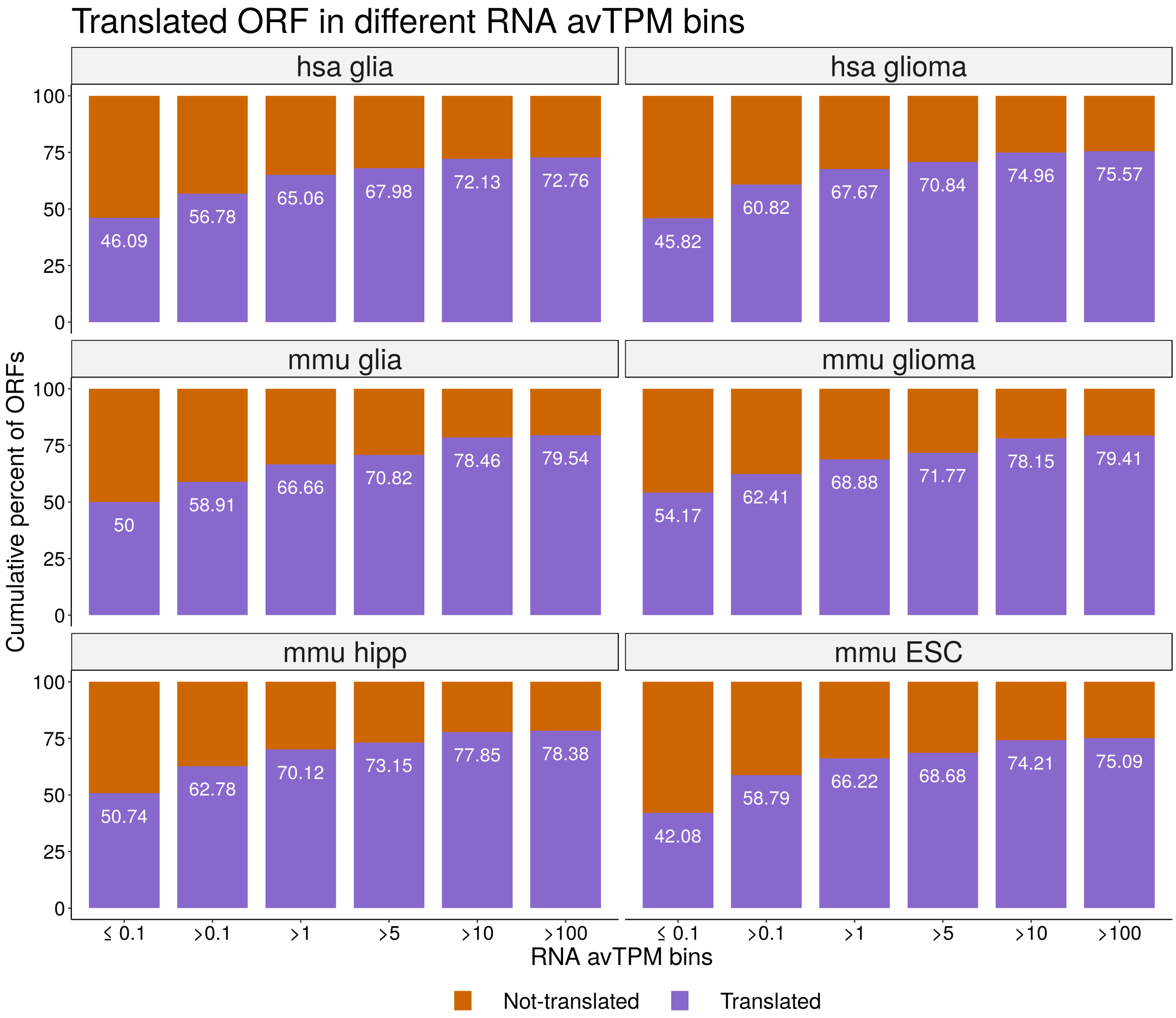


**Supplementary Figure 2.** Cumulative percentage of ORFs predicted as translated (purple) and not translated (red) by ORQAS, as a function of the cut-off for the RNA-seq abundance value (TPM) (averaged across replicates) (x axis). Data is shown for human samples of glia (hsa glia) and glioma (hsas glioma) and the mouse samples of glia (mmu glia), glioma (mmu glioma), hippocampus (mmu hipp) and Embrionic Stem Cells (mmu ESC).

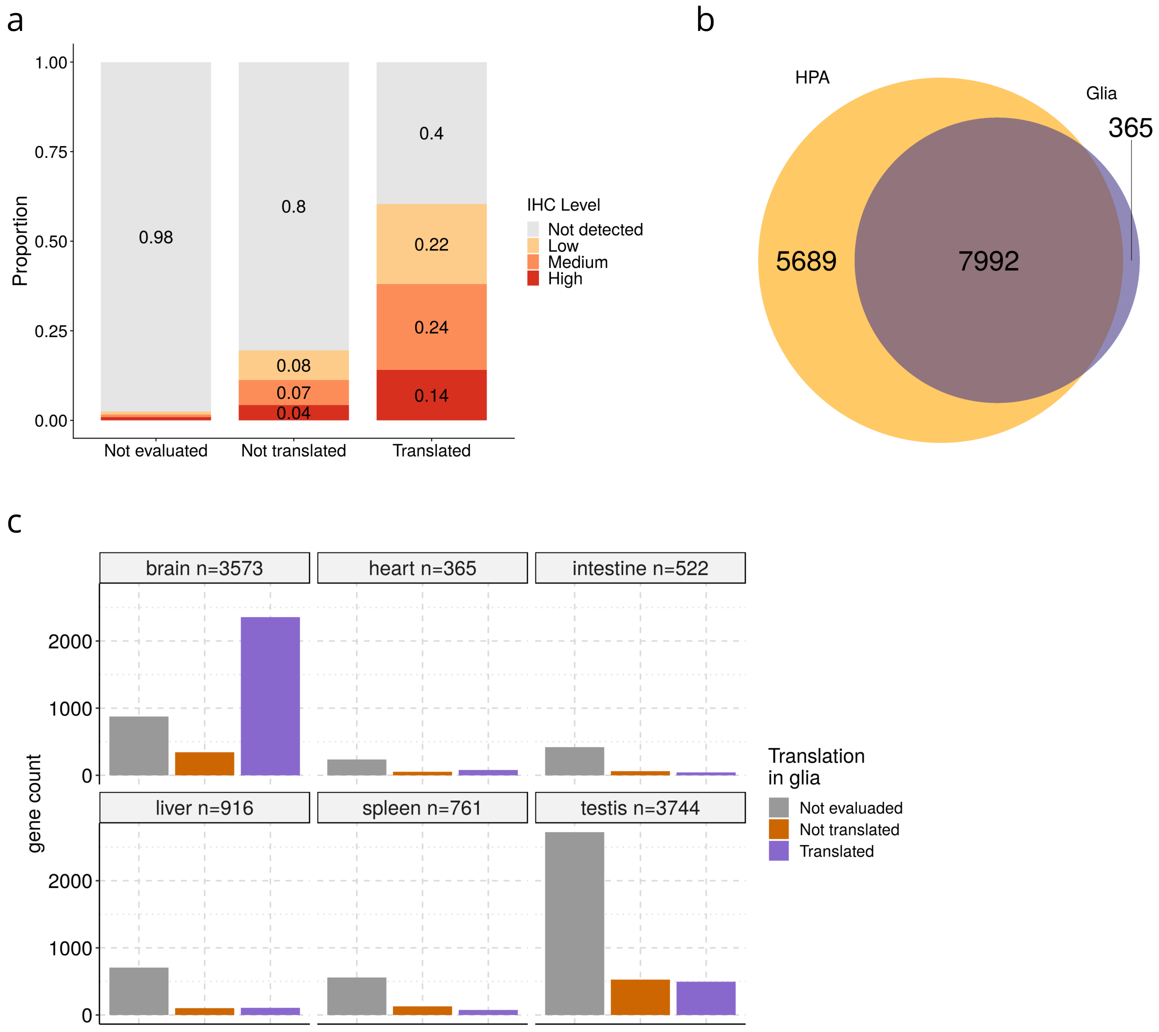


**Supplementary Figure 3. (a)** For the predicted translated isoforms (translated), the cases that did not pass the thresholds of uniformity and periodicity (not-translated), and those without enough read data to be tested (not evaluated) in human glia (hsa glia), the plot shows the proportion of cases for which the gene has evidence from immunohistochemistry (IHC) in cortex, separated as high, low, and medium expression, from the Human Protein Atlas (THPA). We also indicate the cases not detected with IHC (not-detected). Singletons (single-ORF genes) were not included. **(b)** The plot shows the number of genes with predicted translated ORFs with evidence of protein expression in the Human Protein Atlas from a combination of features: Mass Spectrometry, Immunohistochemistry and Uniprot. Translation predictions correspond to human glia Ribo-seq. Singletons were not included. **(c)** For genes with tissue specific RNA and protein expression, as annotated in the Human Protein Atlas, in six different tissues (brain, heart, intestine, liver, spleen and testis), the plot shows ORQAS predictions in the human glia sample (hsa glia) for the candidate translated isoforms (translated), the cases that did not pass the threshold of uniformity and periodicity (not-translated), and those without enough read data to be tested (not evaluated).

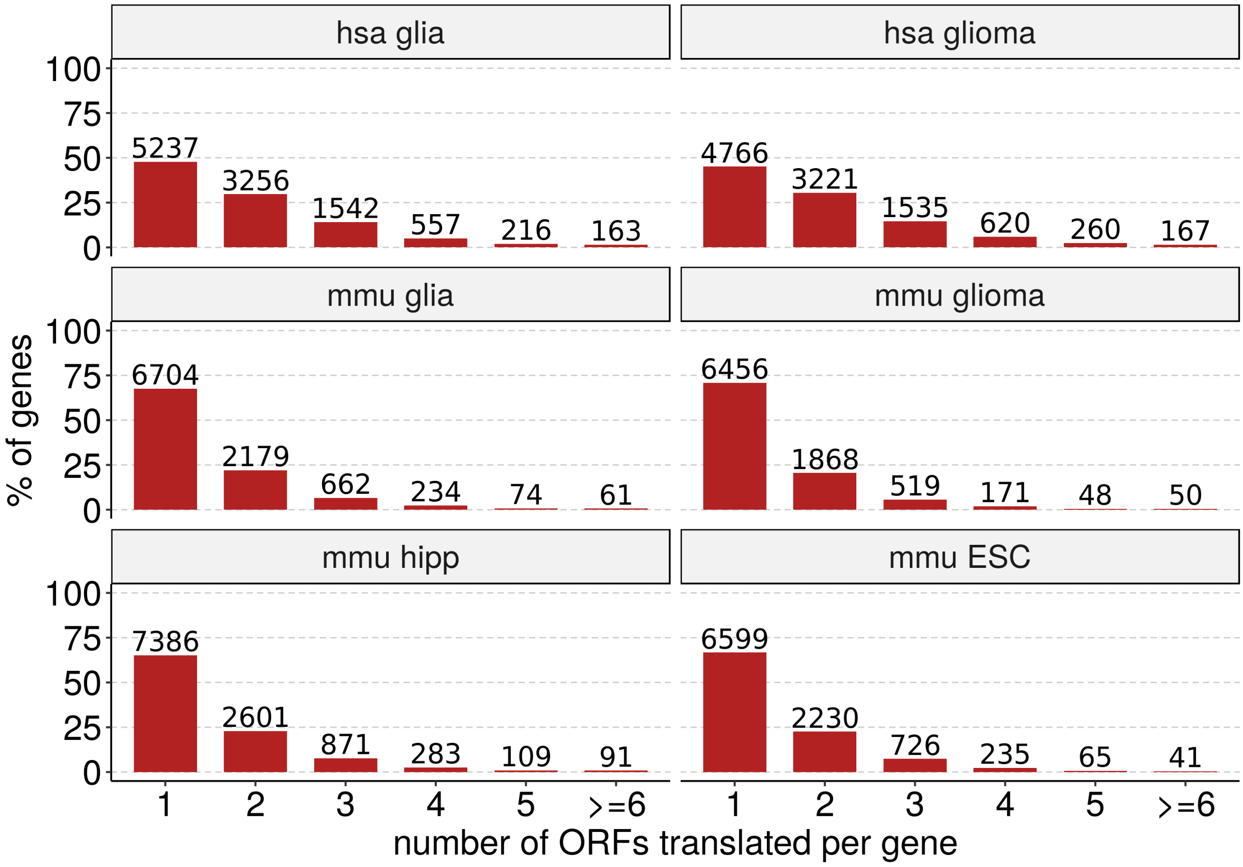


**Supplementary Figure 4.** Distribution of the number of different ORFs translated per gene in the human samples of glia (hsa glia) and glioma (hsa glioma) and mouse samples of glia (mmu glia), glioma (mmu glioma), hippocampus (mmu hipp) and Embrionic Stem Cells (mmu ESC).

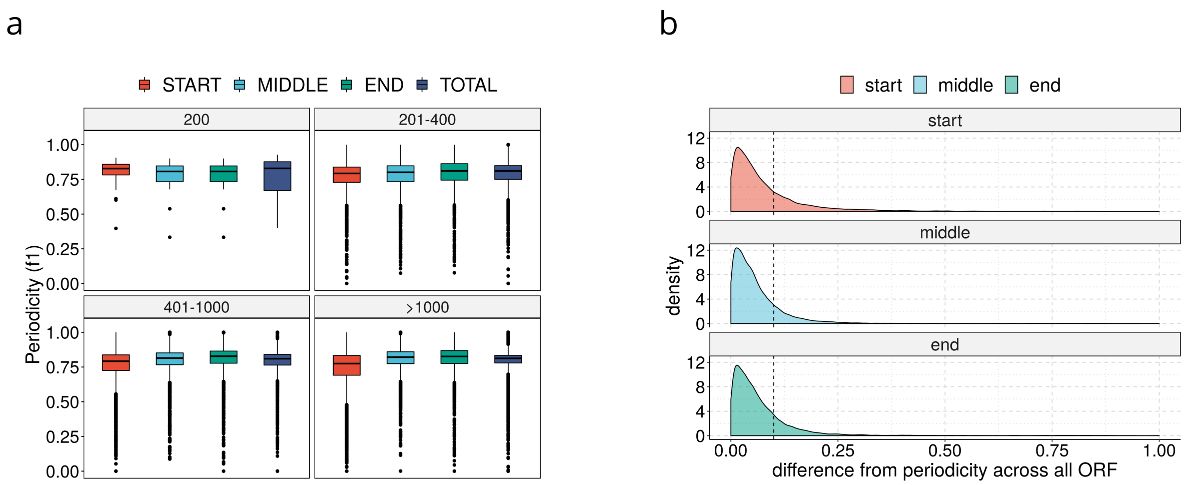


**Supplementary Figure 5. (a)** Distribution of periodicity values calculated for the segments resulting when partitioning the ORFs in three portions of equal length at the start (red), in the middle (light blue) and at the end (green). We include the periodicity value calculated globally for the whole ORF length as it is used by ORQAS (dark blue). ORFs are grouped according to its length (200nt, 201-400nt, 401-1000nt and >1000nt). **(b)** Distribution of the differences of the periodicity values in each of the segments defined in (a) with respect to the total value. The dashed line is set at 0.10.

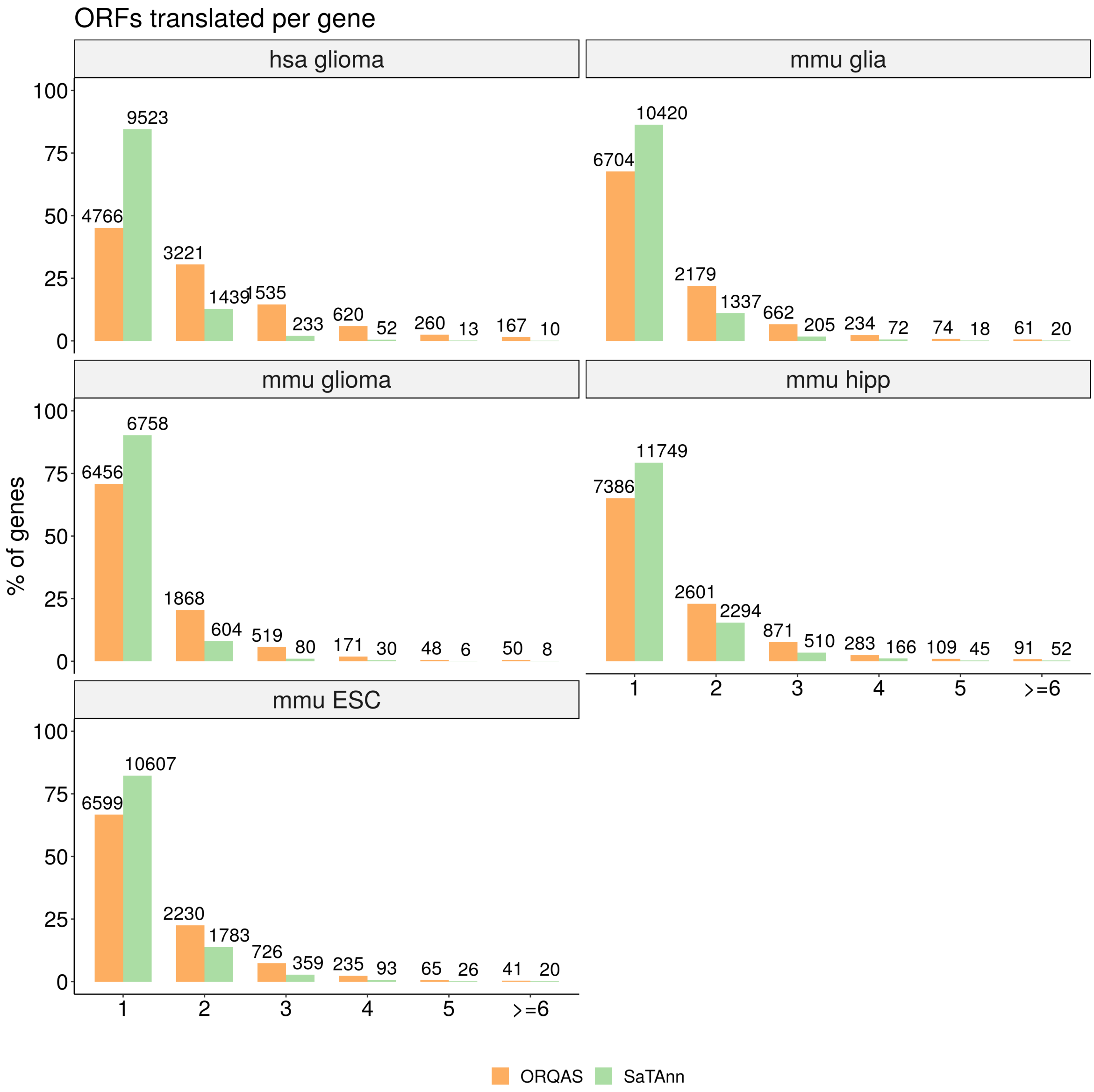


**Supplementary Figure 6.** Distribution of the number of different ORFs translated per gene according to ORQAS (orange) and SaTAnn (green) in the human samples of glia (hsa glia) and glioma (hsa glioma) and mouse samples of glia (mmu glia), glioma (mmu glioma), hippocampus (mmu hipp) and Embrionic Stem Cells (mmu ESC).

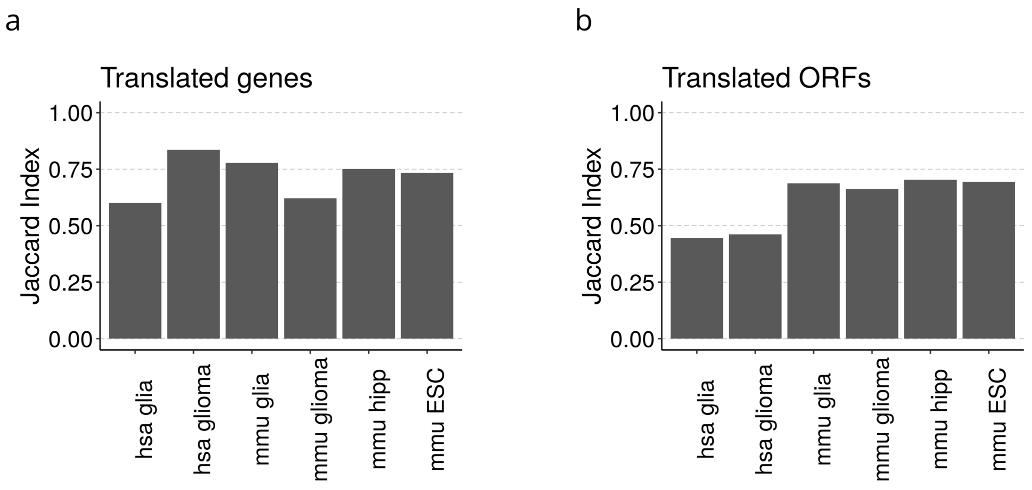


**Supplementary Figure 7. (a)** Overlap between the genes with at least one ORF predicted to be translated by ORQAS and/or by SaTAnn calculated as a Jaccard Index (number of genes with at least one ORF predicted to be translated by both methods divided by the total number with genes with at least one ORF predicted to be translated by any of the methods). **(b)** Overlap between ORF predicted to be translated in ORQAS and in SaTAnn calculated as a Jaccard Index (number of ORFs predicted to be translated by both methods divided by the total number ORFs predicted to be translated in any of the methods).

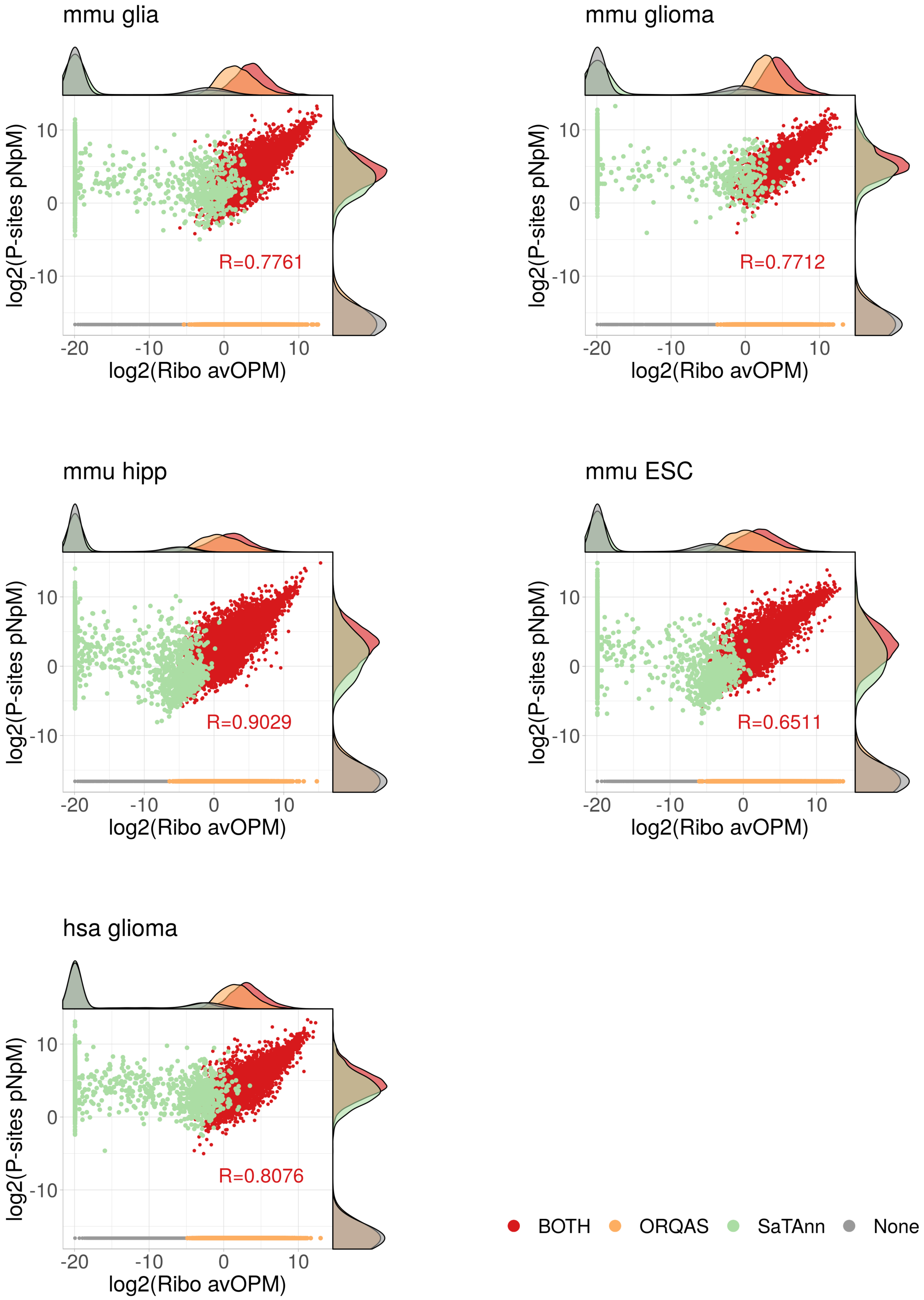


**Supplementary Figure 8.** Correlation between the average abundance in Ribosome space measured as ORFs per Million (OPM) by ORQAS and as P-sites per Nucleotide per Million (P-sites pNpM) by SaTAnn for the samples of human glioma (hsa glioma) and mouse samples of glia (mmu glia), glioma (mmu glioma), hippocampus (mmu hipp) and Embrionic Stem Cells (mmu ESC).

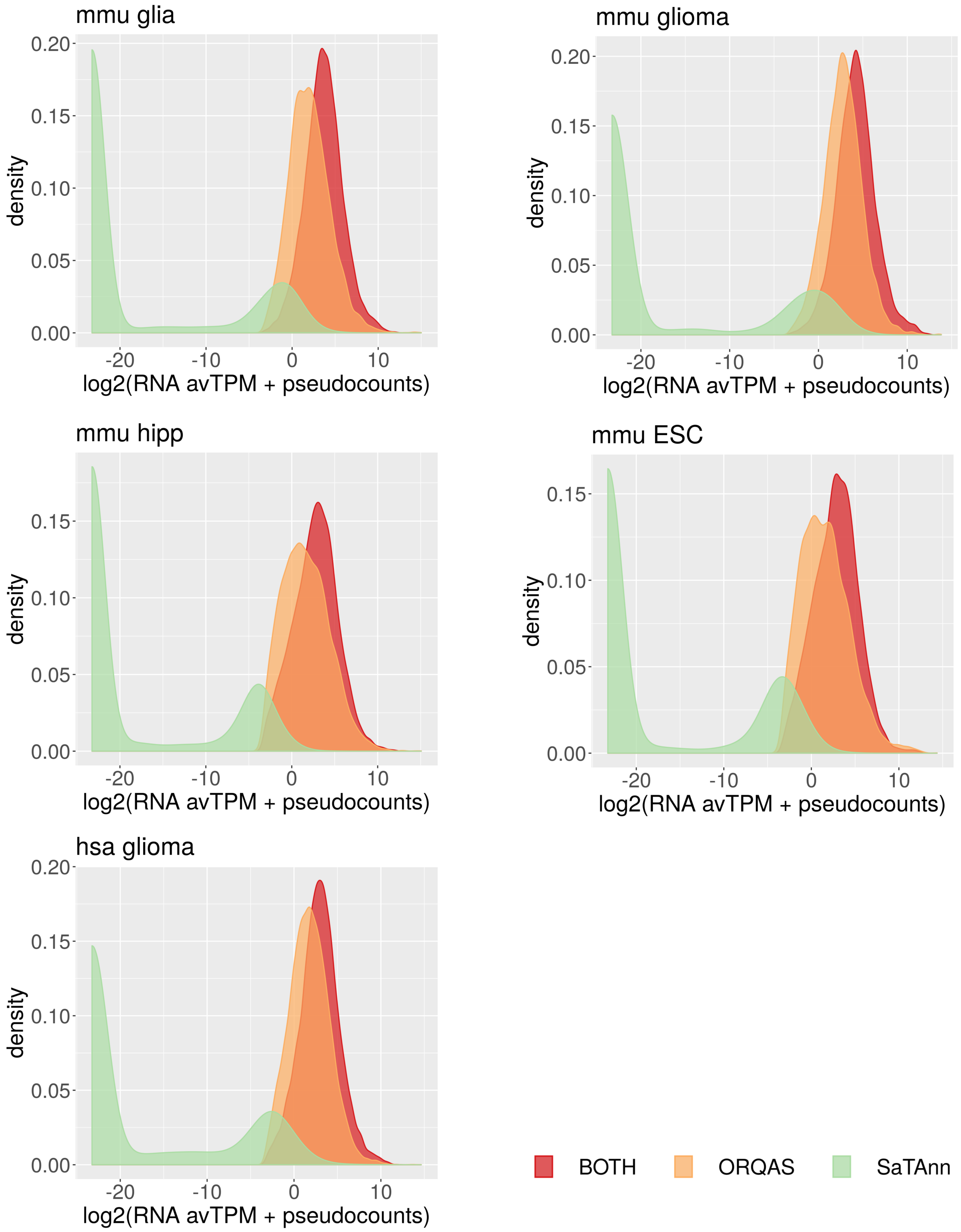


**Supplementary Figure 9.** Distribution of the average RNA expression measured in TPM (averaged over two replicates) for ORFs that are predicted to be translated only by ORQAS (orange), only by SaTAnn (green) or by both methods (red) in human glioma (hsa glioma), mouse glia (mmu glia), mouse glioma (mmu glioma), mouse hippocampus (mmu hipp) and mouse Embrionic Stem Cells (mmu ESC).

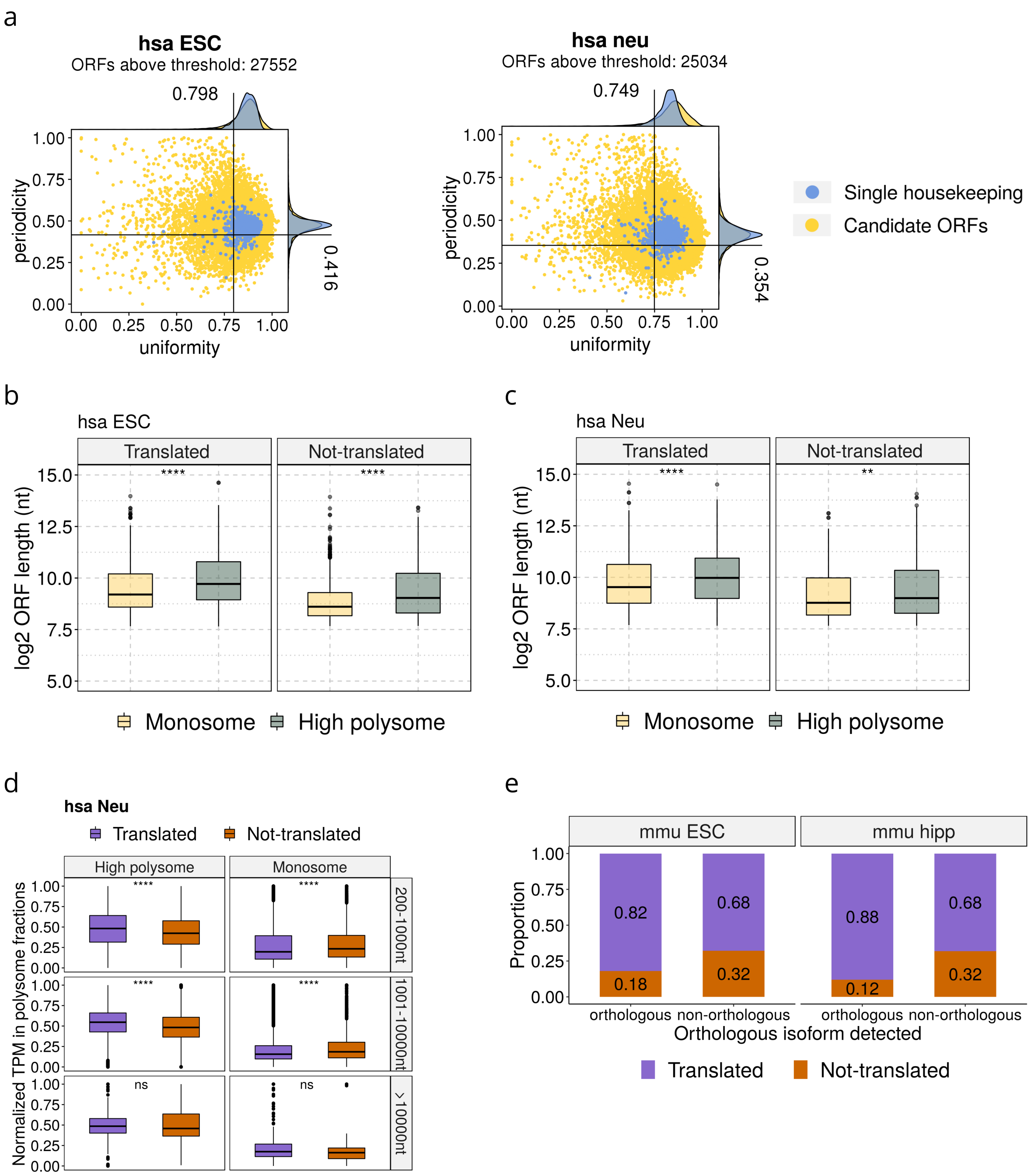


**Supplementary Figure 10. (a)** Uniformity (x axis) versus periodicity (y axis) for the ORFs with RNA expression TPM > 0.1 and at least 10 Ribo-seq reads assigned. In blue we indicate single-ORF genes with protein expression evidence in 37 tissues from THPA, and in yellow the rest of ORFs considered. Uniformity is measured as the percentage of maximum entropy and periodicity is measured in the first annotated frame. We show the data for the samples of human ESCs (left panel) and human differentiated neurons (right panel). **(b)** Distribution of the length in nucleotides of the ORFs expressed (normalized TPM > 0) only in the high polysome (left panels) and monosome (right panels) fractions of human Embrionic Stem Cells (has ESC) for translated isoforms (p-value 2.5e-05) and for isoforms with RNA expression (TPM>0.1) but predicted as not translated (p-value 0.0463). **(c)** Distribution of the length in nucleotides of the ORFs expressed (normalized TPM > 0) only in the high polysome (left panels) and monosome (right panels) fractions of human Neural cells (has Neu) for translated isoforms (p-value 4.14e-15) and for isoforms with RNA expression (TPM>0.1) but predicted as not translated (p-value 0.0028). **(d)** We show the distribution of the relative abundance in high polysome (left panels) and monosome (right panels) fractions of human Neural cells (has Neu) for translated isoforms and for isoforms with RNA expression (TPM>0.1) but predicted as not translated. The plot shows the results for three different ORF lengths: 200-1000nt (high polysome p-value 6.35e-25 and monosome p-value 1.05e-08), 1001-10000nt (high polysome p-value 4.1e-53 and monosome p-value 2.9e-10) and longer than 10000nt (high polysome p-value 0.62 and monosome p-vaue 0.53). **(e)** For the set of ORFs encoding a human-mouse orthologous protein pair (orthologous) and for those encoding proteins without an orthologous pair in mouse (non-orthologous) we plot the percentage that are predicted to be translated (translated) and the ones that did not pass the uniformity and periodicity thresholds (not-translated). We show here the results for mouse ESCs (p-value = 5.228e-124 Fisher test) and for mouse hippocampus (p-value = 9.821e-311).

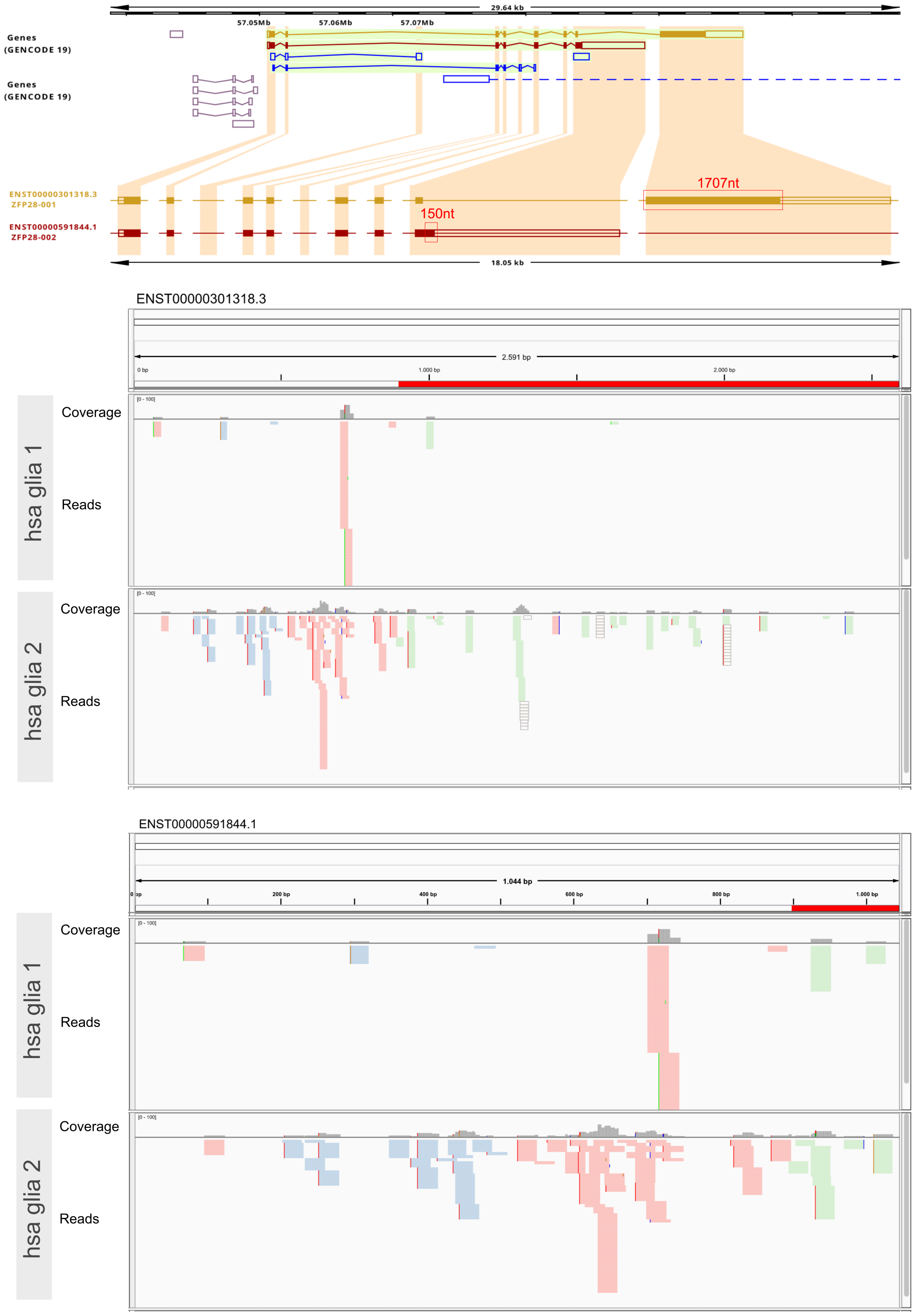


**Supplementary Figure 11.** The upper panel shows the Ensembl annotation of the transcripts of *ZFP28* gene (ENSG00000196867) highlighting in red boxes the specific-sequence regions. Lower panels depict Ribo-seq reads in the human glia samples (hsa glia) mapping to the CDS of two isoforms of this gene ENST00000301318 (top) and ENST00000591844 (bottom), where specific-sequence regions are represented by the region highlighted in red. For each CDS we show the coverage of reads. The reads are colored according to the number of regions in the genome where the reads map: green (one), blue (two) and pink (three).

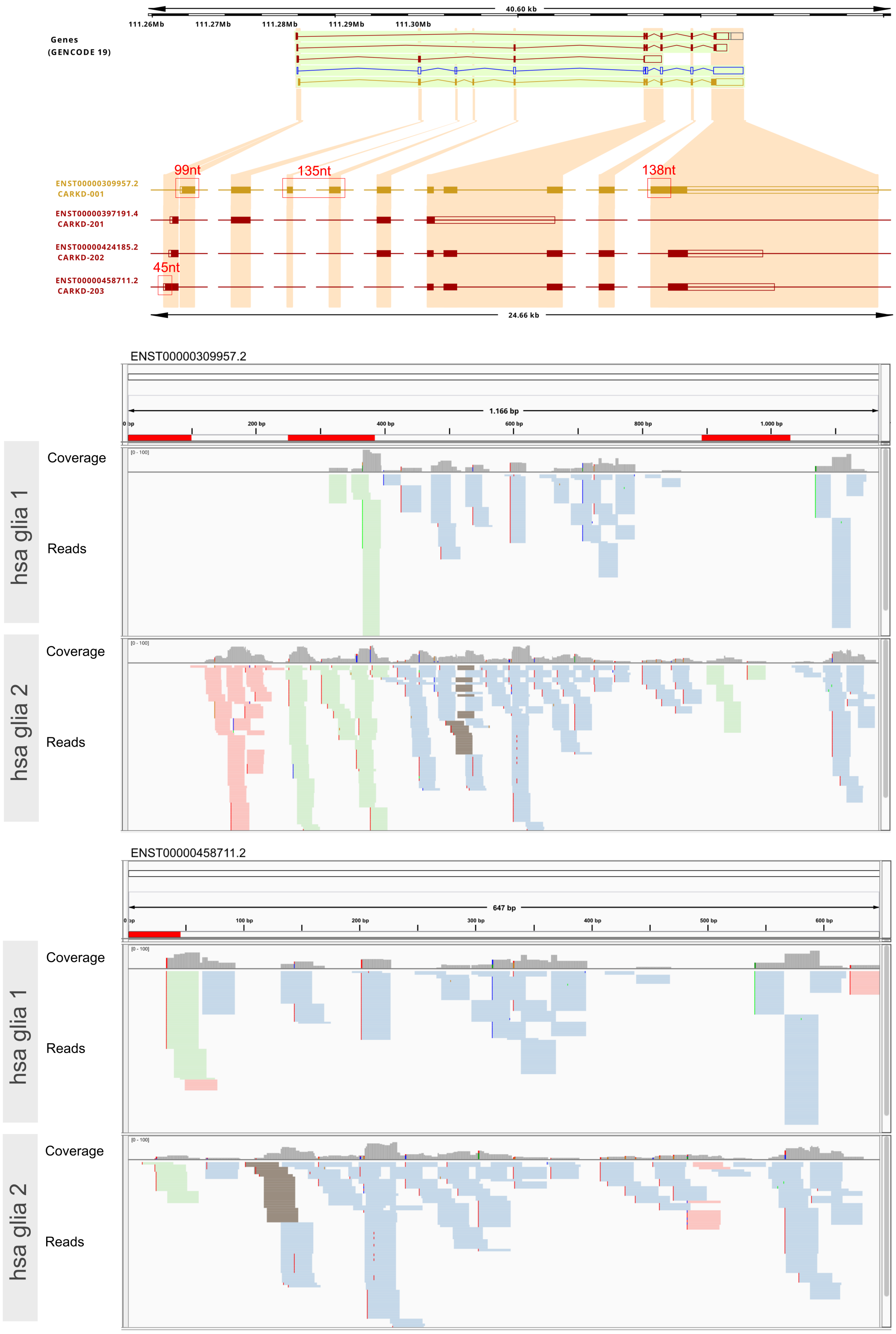


**Supplementary Figure 12.** The upper panel shows the Ensembl annotation of the transcripts of NAXD gene (ENSG00000213995) highlighting in red boxes specific-sequence regions. Lower pannels depict Ribo-seq reads in the human glia samples (hsa glia) mapping to the CDS of two isoforms of this gene ENST00000309957 (top) and ENST00000458711 (bottom), where specific-sequence regions are represented by the region highlighted in red. For each CDS we show the coverage of reads. The reads are colored according to the number of regions in the genome where the reads map: green (one), blue (two) and pink (three).

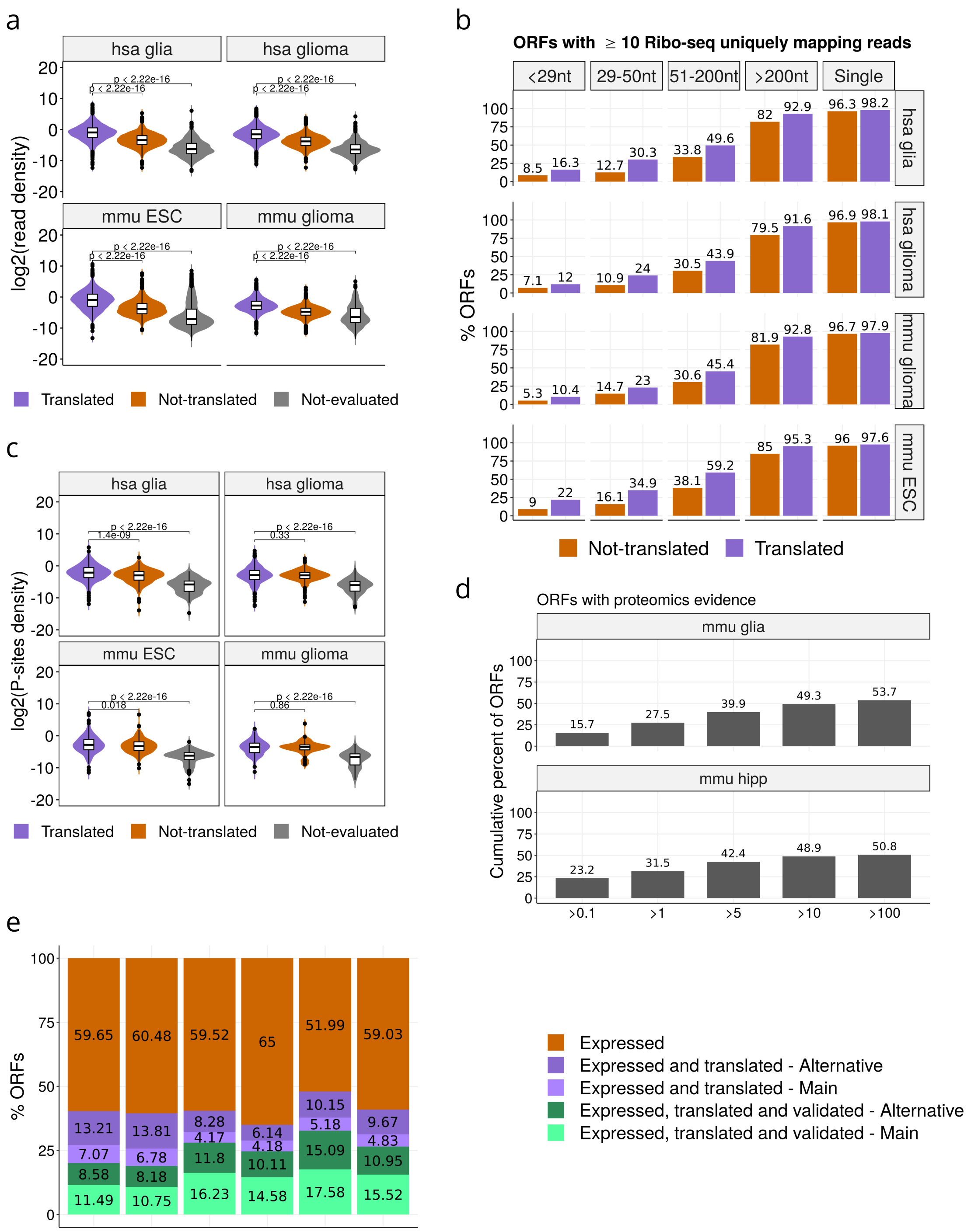


**Supplementary Figure 13. (a)** For the human samples of glia (hsa glia) and glioma (hsa glioma) and the mouse samples of glioma (mmu glioma) and Embrionic Stem Cells (mmu ESC) we show the density of Ribo-seq reads per nucleotide over the isoform-specific sequence regions, calculated as the uniquely mapping read-count over region length in log2 scale for isoform-specific sequences. Distributions are given for predicted translated isoforms, for isoforms that did not pass the threshold of uniformity and periodicity (not translated), and for the isoforms with low expression (TPM<0.1) (not evaluated). **(b)** For the human samples of glia hsa glia) and glioma (hsa glioma) and the mouse samples of glioma (mmu glioma) and Embrionic Stem Cells (mmu ESC) the plot shows the percentage of regions with at least 10 uniquely mapping Ribo-seq reads in isoform-specific sequences over the total number of isoforms with an isoform-specific sequence defined according to the length of the region. **(c)** For the human samples of glia hsa glia) and glioma (hsa glioma) and the mouse samples of glioma (mmu glioma) and Embrionic Stem Cells (mmu ESC) we show the density of Ribo-seq reads per nucleotide over the isoform-specific ORFs, calculated as the counts per nucleotide based on the estimated P-site positions over region length in log2 scale for isoform-specific ORFs. **(d)** For the mouse samples of glia (mmu glia) and hippocampus (mmu hipp), the plot shows the overall cumulative percentage of ORF-specific regions with 1 or more mass-spectrometry peptides according to increasing cut-offs of average RNA-seq abundance values (TPM). The plot shows the combined results for both types of regions: isoform-specific sequence regions and isoform-specific ORFs for isoforms predicted to be translated. **(e)** For each sample, and for all ORFs with sufficient RNA-seq expression (TPM > 0.1), we show the proportion of main isoforms and alternative isoforms predicted to be translated from Ribo-seq reads and the proportion that were validated. These plots do not include the genes with a single protein-coding isoform.

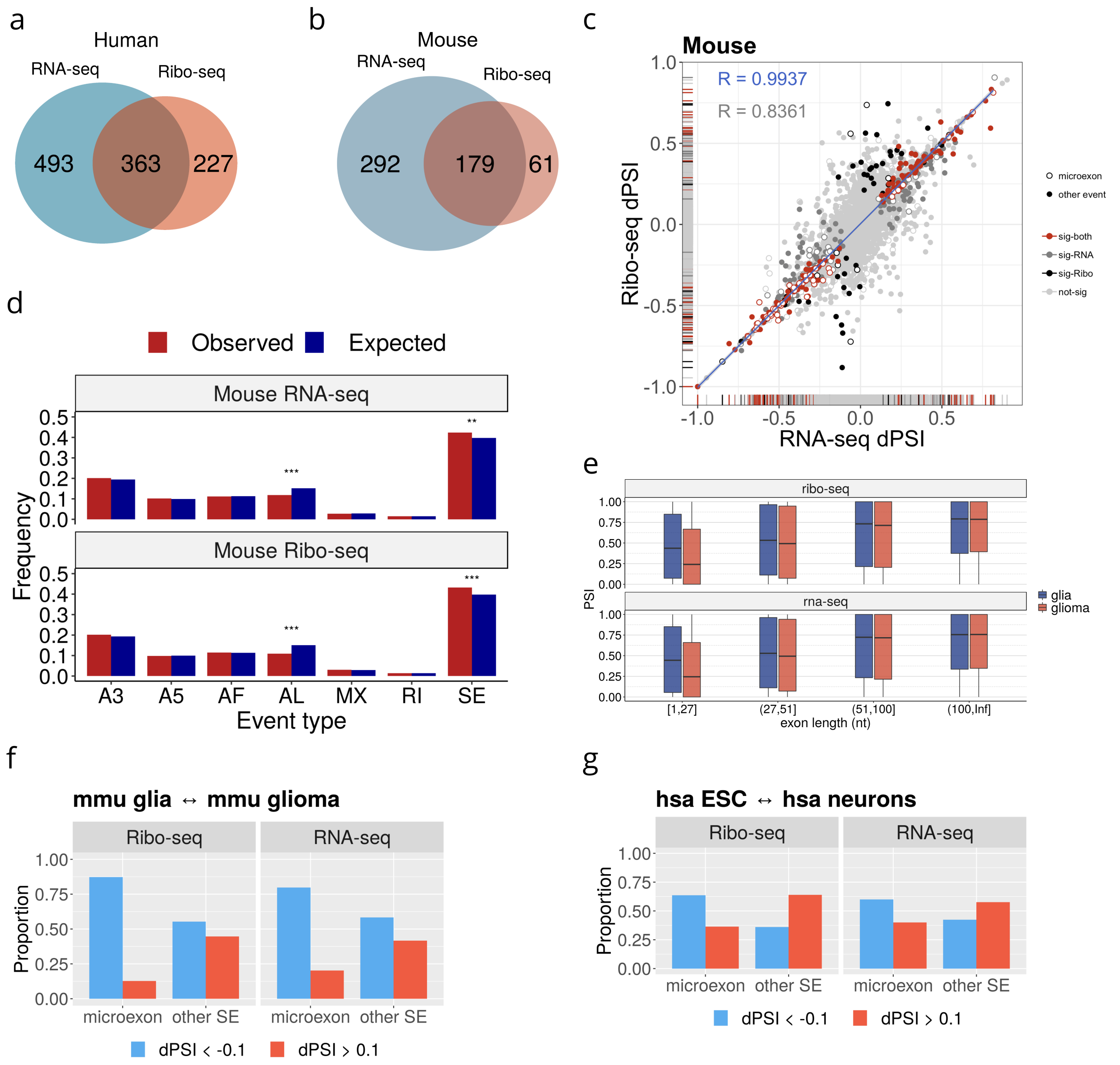


**Supplementary Figure 14. (a)** Overlap of events changing significantly (dPSI > 0.1 and p-value < 0.05) with RNA-seq and with Ribo-seq for human. **(b)** Overlap of events changing significantly (dPSI > 0.1 and p-value < 0.05) with RNA-seq and with Ribo-seq for mouse. **(c)** Correlation of changes in splicing and translation in events in mouse. **(d)** Proportions of events calculated in RNA space (upper panel) or Ribosome space (lower panel). In blue we show the proportion of alternative splicing events calculated with SUPPA that overlap coding regions in mouse, whereas in red we show the events that show a significant change using RNA-seq comparing mouse glia and glioma. Even types are alternative 3'ss (A3) and 5'ss (A5), alternative first (AF) and last (AL) exon, mutually exclusive (MX) exon, retained introns (RI) and skipping exon (SE). There is significant enrichment of SE events for RNA (Fisher's test p-value = 3.22e-03) and Ribo-seq (p-value = 3.04e-04); and significant depletion of AL events for RNA (2.17e-13) and Ribo-seq (2.86e-18). **(e)** Inclusion level in PSI values of skipping exon (SE) events according to the length of the alternative exon in nucleotides (nt) for the human glia and glioma samples. Microexons were defined as exons of length <52nt. **(f)** Enrichment of microexons with an impact in RNA splicing and ORF translation in mouse from the comparison of glia and glioma samples. In the figure, dPSI is used to indicate the difference in relative abundance in both RNA and Ribosome spaces. **(g)** Difference in high polysome fraction, measured as dPSI, between human neuronal samples and ESCs (y axis) for microexons with a significant change in Ribosome space. As before, dPSI indicates the difference in relative abundance in Ribosome space.

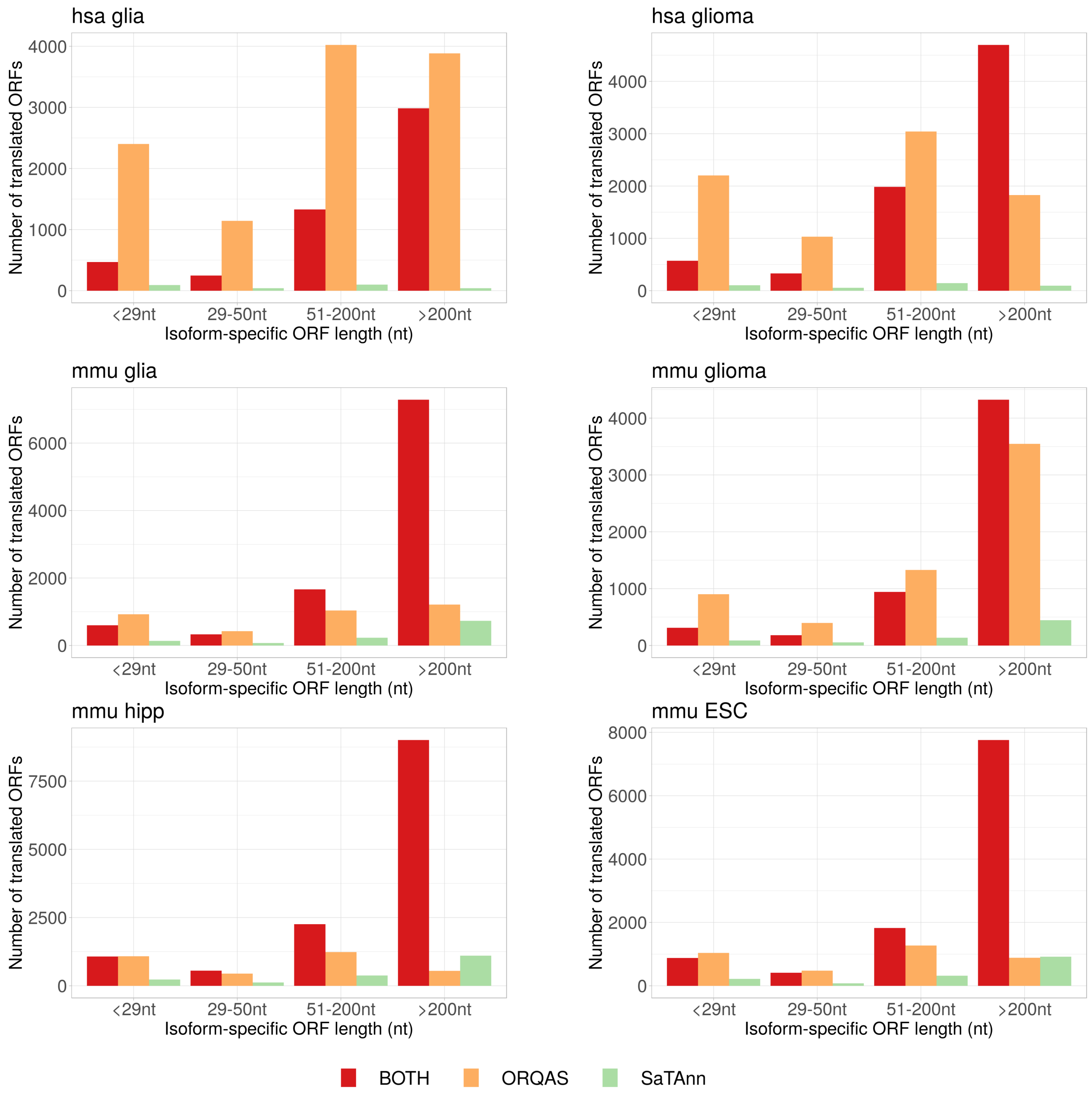


**Supplementary Figure 15.** Number of isoforms containing isoform-specific ORF regions separated by region length and according to whether they were predicted to be translated only by ORQAS (orange), only by SaTAnn (green) or by both methods (red). Data is shown for the samples of human glioma (hsa glioma) and mouse samples of glia (mmu glia), glioma (mmu glioma), hippocampus (mmu hipp) and Embrionic Stem Cells (mmu ESC).

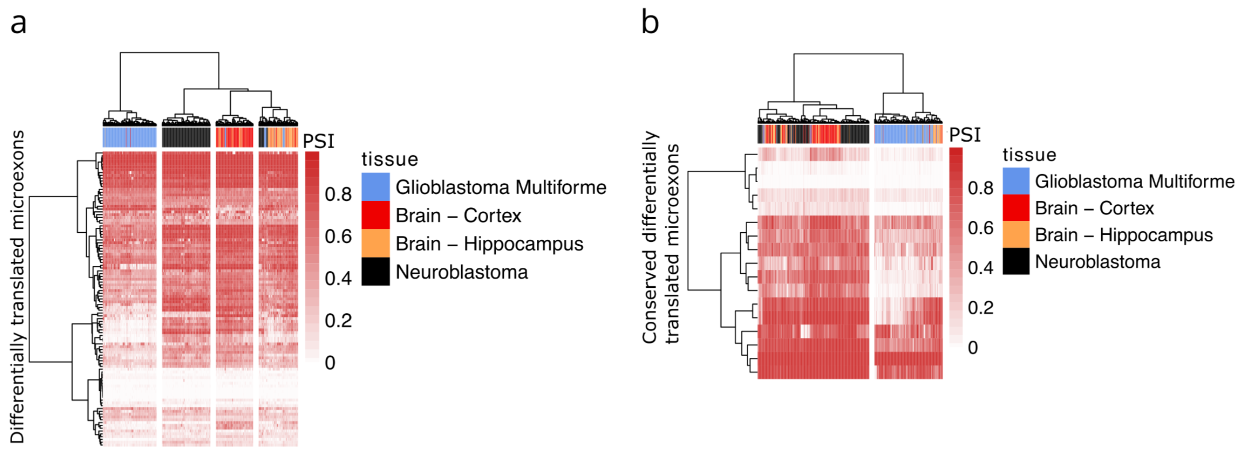


**Supplementary Figure 16. (a)** Patterns of inclusion in normal brain and hippocampus samples from GTEX, gliblastoma multiforme from TCGA and neuroblastoma from TARGET, for microexons that were differentially translated between glia and glioma. The heatmap shows the Percent Spliced In (PSI) values for each sample. **(b)** Patterns of inclusion in normal brain and hippocampus samples from GTEX, gliblastoma multiforme (GBM) from TCGA and neuroblastoma (NB) from TARGET, for microexons that were differentially translated between glia and glioma and this pattern was conserved in human and mouse. The heatmap shows the Percent Spliced In (PSI) values for each sample.
